# AVJ16 inhibits the RNA binding protein IGF2BP1 in lung adenocarcinomas and prevents tumor growth in mice

**DOI:** 10.1101/2024.08.15.608038

**Authors:** Nadav Wallis, Tehila Gershon, Sojod Shaaby, Froma Oberman, Myriam Grunewald, Deborah Duran, Amandeep Singh, Gilad Vainer, Vladimir S. Spiegelman, Arun K. Sharma, Joel K. Yisraeli

**Author notes:** Nadav Wallis, Department of Cellular and Molecular Medicine, University of California San Diego, La Jolla, California, USA.

## Abstract

IGF2BP1 is an oncofoetal RNA binding protein that is expressed in many tumors. We have recently described a small molecule inhibitor of IGF2BP1, termed AVJ16, that prevents binding of the protein to its RNA targets by directly associating with the protein. Here, using a multi-omics approach, we have analyzed the effects of this inhibition on RNA binding, RNA expression, and protein expression. AVJ16 treatment downregulates RNAs encoding members of several pro-oncogenic signaling pathways, including Hedgehog, Wnt, and PI3K-Akt, and there is a strong correlation between IGF2BP1 RNA binding, RNA expression, and protein expression. AVJ16 treatment of lung adenocarcinoma (LUAD) cells in culture causes a strong reduction in proliferation, colony formation, invasion, and spheroid growth while enhancing apoptosis and cell death. All of these effects are limited to cells expressing IGF2BP1. LUAD cells treated with AVJ16 show a pronounced reduction in vital dye efflux, often correlated with enhanced chemosensitivity. In syngeneic LUAD xenografts in mice, IP injection of AVJ16 prevents tumor growth, and incubation with AVJ16 induces cell death in human organoids derived from IGF2BP1-expressing LUADs but not from healthy lung tissue. These results suggest that AVJ16 is a promising candidate for mono- and/or adjuvant therapy directed against tumors expressing IGF2BP1.

## INTRODUCTION

The IGF2BP RNA binding protein family consists of three paralogs, IGF2BP1, IGF2BP2, and IGF2BP3, that regulate stability, intracellular localization, translation, and/or alternative splicing of their mRNA targets.^1^ All three paralogs are expressed during embryogenesis in a variety of tissues, but IGF2BP1 and 3 are downregulated postnatally. In many types of cancers, however, the expression of one or more of the paralogs is upregulated.^2, 3^ We have focused on the involvement of IGF2BP1 in lung adenocarcinoma (LUAD), where we have observed that IGF2BP1 synergizes with mutant Kras to accelerate tumor progression.^4^ In general, IGF2BP1’s expression in numerous types of cancers is associated with upregulation of key pro-oncogenic RNAs, poor prognosis, and reduced survival.^5^ Studies using mouse models in which IGF2BP1 was knocked out or knocked down^6–12^ suggest that molecules inhibiting IGF2BP1 could have therapeutic potential in cancer treatment. We previously isolated a small molecule termed 7773 that interacts with a hydrophobic surface at the boundary of IGF2BP1 KH3 and KH4 domains, inhibiting its RNA binding activity^13^. Optimization of 7773 led to development of a related molecule, AVJ16, with an improved affinity for IGF2BP1 (K_d_=1.4μM). AVJ16 demonstrates the same specificity as 7773 for IGF2BP1, does not bind to IGF2BP2/3 in vitro, has no effect on cell lines that express little to no IGF2BP1, and is particularly efficient at preventing wound healing and cell proliferation and migration^14^.

IGF2BP1 is known to target a wide range of RNAs involved in diverse cellular processes, although the exact set of targets varies depending on the specific cell type and physiological context. Several high-throughput sequencing-based techniques have revealed hundreds to thousands of potential targets for this protein under different experimental conditions and analysis methods^15–17^. Among the many IGF2BP1 mRNA targets are several oncogenes including LIN28B and HMGA2^18^, Bcl2^17^, KRAS^19^, E2F1^9^, Cmyc^20^, CD44^21^, GLI1^22^, BTRC^23^, and CTNNB1^24^. IGF2BP1 binding is mediated by specific cis-acting sequence and structural motifs predominately located within the 3’ untranslated region (UTR) of target mRNAs, although binding is also observed, to a lesser degree, in coding domain sequences (CDS) and other regions in the RNA.^17^ IGF2BP1, like its paralogs IGF2BP2 and 3, is also an RNA N^6^-methyladenosine (m^6^A) reader^25^ and recently has been identified as a part of a complex that helps facilitate nuclear export of circular RNA.^26^ By positively regulating a large number of pro-oncogenic, anti-apoptotic, and chemoresistance RNAs, IGF2BP1 plays important roles in many diverse pathways that promote tumor progression.^27^ Consequently, inhibition of IGF2BP1 can exert a pleiotropic effect on many RNAs and pathways simultaneously.

Here, we characterize the mechanisms of action of AVJ16 as an IGF2BP1 inhibitor by assessing the effects of AVJ16 treatment in LUAD. We have performed a multi-omic analysis to identify changes in direct RNA binding sites (eCLIP), RNA expression (RNAseq), and protein expression (Mass spectrometry), thereby revealing the global effect of the inhibitor on cancer progression, and have identified novel mechanisms through which IGF2BP1 influences LUAD. In a syngeneic mouse xenograft model, AVJ16 almost completely inhibits growth of the primary tumor. These results suggest that AVJ16 holds promise as a novel therapeutic approach for treating lung, and potentially other, IGF2BP1-expressing tumors.

## MATERIALS AND METHODS

### Tissue culture

H1299 cells were maintained in RPMI medium, and LKR-M and RKO cells were maintained in DMEM (Biological Industries - Israel). Both media contained 10% FCS (Biological Industries) and 10 μg/mL ciprofloxacin (Bayer).

### Cell migration assay

H1299 and HEK293 cells were seeded in 96 IncuCyte® ImageLock plates (20X10^3^ cells per well) for 24 hours, to near confluency (95%), before the addition of increasing concentrations of compound. After a further 24 hours, wells were scratched using the IncuCyte® 96-well WoundMaker Tool, and the cells cultured for an additional 48 hours using the IncuCyte® S3 Live-Cell Analysis System (Essen BioScience). The plate was imaged at increments of 120 minutes for a period of 48 hours, and then analyzed for relative wound healing.

### LDH cytotoxicity assays

LDH Cytotoxicity Assay is a colorimetric assay that assesses Lactate dehydrogenase (LDH) release to the media. The protocol was performed as described by the manufacture (Invitrogen). 5000 cells were cultured for 1day in 96-well plate and after 24 hours, compound/DMSO was added for 96 hours. All measurements were done using a plate reader (Infinite M200 PRO NanoQuant, Tecan). Optical density was proportional to the number of cells.

### Apoptosis assays

For the IncuCyte apoptosis assay, to determine the effect of AVJ16 on apoptosis in cells, we utilized the Caspase-3/7 apoptosis assay and monitored the experiment by IncuCyte live cell imaging system. Briefly, H1299 and LKRM cells were seeded at 5000 cells per well in a 96-well plate (Corning). After incubating overnight, 4μM AVJ16 and the Caspase-3/7 reagent (Sartorius) were added. The plate was imaged at increments of 120 minutes for a period of at least 48 hours. The analysis was done with IncuCyte® S3 Live-Cell Analysis System (Essen BioScience).

### FACS

The apoptosis assay was performed using FACS using Annexin V–CF Blue/7-amino-actinomycin D (7-AAD) Apoptosis Detection Kit (ab214663, Abcam) according to the manufacturer’s protocol. Briefly, cells were detached using 0.05% trypsin and washed twice with PBS. Then, samples were resuspended in 1× annexin-binding buffer and incubated with Annexin V-FITC and 7-AAD for 15 min at 37 °C, avoiding light. The stained samples were analyzed on a BD LSR II analyzer Flowcytometer at an excitation wavelength of 488 nm and emission filters of 525 and 625 nm. Acquisition was done with BD FACSDiva™ software and FCS files was done with FCS-express softwares

### CETSA

H1299 cells were cultured on plates at a density of 1X10^6^ cells per plate, and allowed to adhere overnight. Cells were treated with 10µM AVJ16 or DMSO for three hours. Cells were trypsinized, heated in a thermal gradient between 37°C to 58°C for 3 minutes, and then returned to the 37°C incubator. After the heat shock treatment, cells were lysed in PBS buffer in three cycles of freeze-thaw. Lysates were centrifuged at 13,000 rpm for 10 minutes at 4°C. The supernatant was collected and stored at -80°C. Supernatants were electrophoresed on a 10% SDS-polyacrylamide gel, transferred to nitrocellulose filters, and visualized using antibodies specific to IGF2BP1,2, or 3. Band intensities were quantified using ImageJ software. The melting temperature (Tm) was calculated as the inflection point of the CETSA curve using GraphPad Prism software. All experiments were performed in triplicate.

### Spheroid formation

Cells were seeded in Round-bottom 96-well plates BIOFLOAT™ 96-well Cell Culture Plate with Ultra-low attachment surface (faCellitate) with 2500 cells per well, with or without Matrigel. Cells were centrifuged at 200g for five minutes, and AVJ16 or DMSO was added. Plates were incubated at 37 °C for 8-10 days to allow for spheroid formation. Plates were imaged twice a day and analysis was done with IncuCyte® S3 Live-Cell Analysis System (Essen BioScience).

### Western blot analysis

Western blots were performed as described^4^. For fluorescent westerns, cell lysates were prepared with phosphatase inhibitors (β-Glycerophosphate and Sodium orthovanadate) and a protease inhibitor (cOmplete™, Mini Protease Inhibitor Cocktail, Roche 04693124001), membranes were read using a Li-Core Odyssey laser scanner, and results analyzed using Image Studio Lite software.

The following primary antibodies were used: mouse anti-total ERK (p44/42 MAPK (Erk1/2) L34F12 Cell Signaling 4696s), mouse anti-Kras (CPTC-KRAS4B-2, DSHB) rabbit anti-dpERK, (p44/42 MAPK, Cell Signaling 4370), rabbit anti-alpha/beta tubulin (Cell Signaling 2148), goat anti-rabbit IgG-HRP (Jackson), goat anti-mouse IgG-HRP (Jackson). Secondary antibodies for fluorescent westerns: donkey anti-mouse 800 (Rockland 610-732-124), goat anti-rabbit 680 (Molecular Probes A21076).

### qPCR

Primers were calibrated for several IGF2BP1 RNA targets:

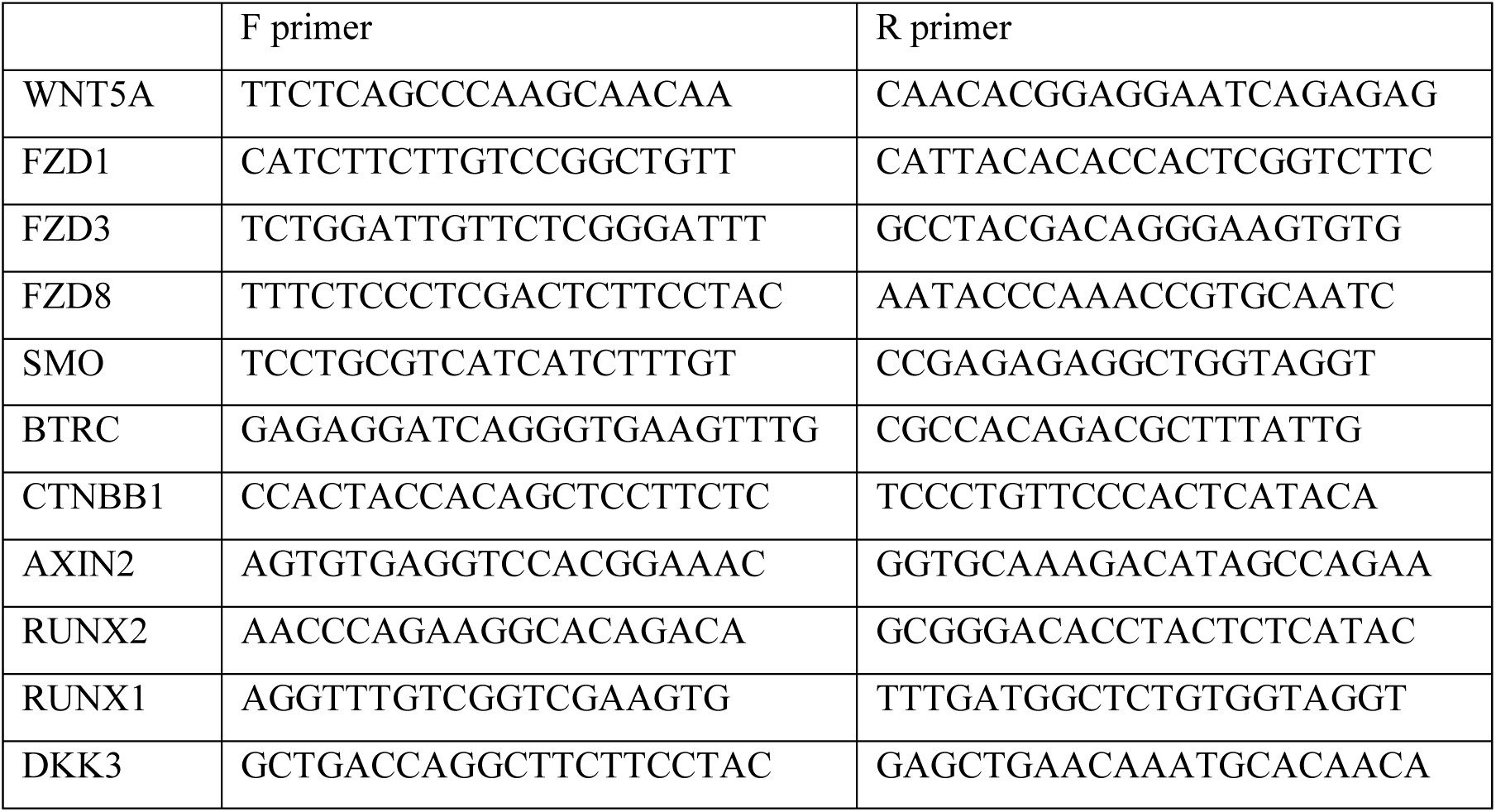

Cells were grown for 12 hours in 12 well plates prior to incubation with the compound at different concentrations for an additional 12 or 24 hours. Total RNA was then extracted using EZ-RNA total RNA isolation kit (Biological Industries) or Trizol (Invitrogen) and cDNA prepared from the RNA using the First Strand cDNA Synthesis kit (Quanta bio). Real time PCR was performed with Fast SYBR Green Master Mix (Thermo Fisher Scientific), and cDNA expression analyzed with the Bio-Rad CFX Manager 3.1.

### Soft agar assay

2X10^5^ cells were initially treated for 48 hours with 4 μM AVJ16 or DMSO. Cells were then harvested, suspended in soft agar (0.35% agarose) mixed with cell culture medium, and reseeded onto 6-well plates containing a preformed layer of 0.5% agarose, at a final concentration of 500 cells per well for LKR-M-GFP and LKR-M-FL and 2500 cells per well for H1299 cells. Cells were fed weekly with 1ml of media containing AVJ16 or DMSO. Visible colonies were determined by adding 120 µl of Iodonitrote-trazolium chloride (1:8 dilution).

### Colony formation

2X10^5^ cells were treated for 48 hours with 4μM AVJ16 or DMSO. Cells were then harvested and seeded onto 24-well plates at a density of 40 cells per well. Cells were grown for an additional one to two weeks in media containing the same treatment. Visible colonies were identified by fixation in 100% methanol for 30 minutes, followed by staining with 0.05% crystal violet for 15 minutes at room temperature. Colonies were rinsed with water and quantified.

### Trans-well migration

Cells were treated for 48 hours with 4μM AVJ16 or DMSO, harvested, and seeded onto a 24-well Boyden Chamber plate at a density of 2X10^4^ cells per well. Cells were seeded in 500µl of serum-free media in the upper chamber with 500µl of media containing 10% fetal bovine serum in the lower chamber and incubated for 24 hours at 37°C. Following this incubation period, cells were gently washed with 1x PBS solution, fixed with 500µl of 4% paraformaldehyde for 10 minutes, and washed again with PBS. Cells were fixed and permeabilized with 500µl of 100% methanol for 20 minutes at room temperature. After several rinses with PBS, they were stained with 500µl of 0.05% crystal violet for 30 minutes. Using cotton swabs, the top side of the filter was gently cleared of any residue, allowing the chamber to dry completely. Stained cells were examined under a Nikon SMZ 25 microscope. The NIS elements software was employed to analyze color intensity and cell counting.

### Trans-well invasion

24-well Boyden Chamber plates were coated with a diluted (1:10) Matrigel solution overnight at 37°C. Cells were treated for 48 hours with 4μM AVJ16 or DMSO, harvested, and seeded in the upper chamber at a density of 10^5^ cells per well in 500μl of FBS-free media, and 500μl of media containing 10% FBS was added to the bottom compartment. Cells were incubated for 24 hours at 37°C and then processed as described above (Trans-well migration).

### Side population (SP) analysis

H1299 cells were incubated either in the presence of 4μM AVJ16 or DMSO for 48 hours or in the presence of 50μM Reserpine for 72 hours. Cells were trypsinized and resuspended in DMEM (Dulbecco’s Modified Eagle Medium) with 2% FCS (Fetal Calf Serum) and HEPES buffer (10 mM, pH 8.5). Hoechst 33342 dye was then added at a final concentration of 2.5 μg/ml and incubated at 37°C for 90 minutes with the dye.

After the incubation period, the cells were washed with ice-cold HBSS (Hanks’ Balanced Salt Solution) containing 2% FCS and 10 mM HEPES. They were then centrifuged at 4°C, and the resulting pellet was resuspended in ice-cold HBSS supplemented with 2% FCS and HEPES. Before sorting, the cell suspension was passed through a 40-μm cell strainer to obtain a single-cell suspension. The analysis and sorting of cells were performed using an Aria III UPG Sorter. The data are presented as the mean ± standard deviation (SD). To determine the significance of differences between groups, an unpaired t-test was conducted using GraphPad Software, Inc. The resulting p-values were evaluated, and values less than 0.05 were considered statistically significant, indicated by asterisks.

### LKR-M lung metastasis model

In total, 2 × 10^5^ LKR-M cells per line were injected subcutaneously into the flanks of randomly selected B6/129 mice, without bias to age, or any other criteria ^28^. Primary tumor volume was measured with callipers three times a week. After 12 days AVJ16 was injected every other day, in a 25% MTD. Four to five weeks post-injection, mice were sacrificed, and lungs and tumors were analysed by haematoxylin and eosin (H&E) and immunohistochemistry (IHC). Number of mice in each cohort: GFP, n = 5; FL, n-male = 7 n-female = 7, each cohort was done twice, the first AVJ16 and the second DMSO control. All mouse experiments described were approved by the Hebrew University Institutional Animal Care and Use Committee.

### Immunohistochemistry

Primary antibody staining of 5 μm sections from 4% buffered formaldehyde fixed tissue were rehydrated, incubated in haematoxylin for 10 min, rinsed in ethanol and then in water. Antigen retrieval was performed using citrate buffer pH 6.0 with a pressure cooker (PickCell Laboratories, Agoura Hills, CA). Slides were stained with the following antibodies: Mice Anti-IGF2BP1 (MBL #RN007P), rabbit anti-Ki67 (Thermo Scientific RM 9106-S), followed by secondary fluorescent staining. Images were captured using Nikon Yokogawa W1 Spinning Disk microscope. Labeling index was calculated from DAPI staining, counting percent of positive nuclei/total nuclei, in an image, in three representative slides.

### Patient-derived organoid culture

Non-Small Cell Lung adenocarcinoma as well as adjacent non-malignant tissue biopsies were obtained after written informed consent from patients treated at the Hadassah Hospital (Jerusalem, Israel) according to the Helsinki protocol (HMO-20-0921). PDTOs were generated according to Li et al^29^. Briefly, fresh biopsies were digested into AdDMEM/F12 supplemented with 1x Antibiotic-Antimycotic, 1x GlutaMAX™ Supplement and 10mM Hepes (AdDMEM/F12^+++^) and containing collagenase II (5mg/ml), Dnase I (10ug/ml) and ROCK inhibitor dihydrochloride (10 μM). Lung cells suspensions were washed with AdDMEM/F12^+++^ and centrifuged at 200g for 5 min. Pellets were washed and resuspended in Growth Factor Reduced Matrigel^®^ (Corning). 40μl of the suspension were plated into pre-warmed 24 well plates and solidified at 37^0^C for 20 min. 500μl of feeding medium (prepared as described in ^29^ for lung cancer and as described in ^30^ for non-malignant lung tissue) were added and replaced every 3 days with freshly prepared feeding medium. Once in 2 weeks, organoids were passaged by mechanical dissociation of the Matrigel domes, washing of the organoids that are plated at a 1:2 ratio in Matrigel and fed with feeding medium.

For drug sensitive testing, organoids were passaged, resuspended in feeding medium and laid on a Matrigel-precoated 96-well plate (as described in ^31^). The day after, drugs were added to the feeding medium and viability assays and further analysis were performed.

### Bioinformatics assays

#### eCLIP-seq Experimental

10x10^6^ UV-crosslinked (400mJ/cm2 constant energy) H1299 cells were lysed in iCLIP lysis buffer and sonicated (BioRuptor). Lysate was treated with RNAse I (Ambion) to shear RNA, after which IGF2BP1 protein-RNA complexes were immunoprecipitated using an anti-IGF2BP1 antibody (MBL, #RN007P). In addition to the RBP-IP, a parallel Size-Matched Input (SMInput) library was generated; these samples were not immunoprecipitated with anti-IGF2BP1 antibody but otherwise were treated identically (to aid in the removal of false positives). Stringent washes were performed, during which RNA was dephosphorylated with FastAP (Fermentas) and T4 PNK (NEB). Subsequently, a 3′ RNA adapter was ligated onto the RNA with T4 RNA ligase (NEB). Protein-RNA complexes were run on an SDS-PAGE gel, transferred to nitrocellulose membranes, and RNA was isolated off the membrane identically to standard iCLIP. After precipitation, RNA was reverse transcribed with AffinityScript (Agilent), free primer was removed (ExoSap-IT, Affymetrix), and a 3′ DNA adapter was ligated onto the cDNA product with T4 RNA ligase (NEB). Libraries were then amplified with Q5 PCR mix (NEB). See Van Nostrand, et al. ^32^ for further details regarding standardized eCLIP experimental workflows.

#### eCLIP-seq Read Processing and Cluster Analysis

Briefly, reads were adapter trimmed (cutadapt), mapped against repetitive elements (with repeat-mapping reads discarded), and then mapped to the human genome with STAR. PCR duplicate reads were removed, and the second (paired-end) read was used to perform peak-calling with CLIPper ^33^. Region-level analysis was performed by counting reads overlapping regions annotated in Gencode (v19). Input normalization of peaks was performed by counting reads mapping to CLIPper-identified peaks in eCLIP and paired SMInput datasets, with significance thresholds of p ≤ 0.001and fold-enrichment ≥ 3. see^32^ for software packages used and additional description of processing steps.

To achieve DBPs, we employed the following methodology: first, we took all CLIPper-called clusters from each IP sample and calculate RPM-normalized read counts of each sample overlapping each cluster. Next, we performed a two-way ANOVA test where we looked for differences in type (IP/ input), treatment, and the interaction of both factors based on the RPM values for each cluster. We then filtered clusters for those that had a p-value < 0.05 for the test of interactions between the two factors. To determine the magnitude of change between the different sample conditions (DBPs), we calculated the average RPM for treated IP, treated input, untreated IP, and untreated input and then used these averages to calculate the treated IP vs. input log_2_ fold change and untreated IP vs. input log_2_ fold change for each cluster. Finally, we calculated the difference between the log_2_ fold changes (treated log_2_ fold change - untreated log_2_ fold change). Clusters were filtered for log_2_ fold change difference > 0.5 or log_2_ fold change difference < -0.5. To compare all DBPs we used DeepTools suit in Galaxy, first all DBPs were converted to Bigwig files with 100bp regions. Then regions were scaled to the same size with computMatrix to calculate scores per genome regions. plotHeatmap was used to plot the heatmap with default parameters.

#### RNA-seq

H1299 cells were treated with 1.5µM of AVJ16 or an appropriate DMSO concentration, for 48 hours. RNA isolation was done with Qiagen RNeasy MinElute Spin Columns, according to the manufacturer’s protocol. RNA quality was assessed with tape station (Agilent TapeStation).

#### Data processing

The NextSeq base calls files were converted to fastq files using the bcl2fastq program as default. The processed reads were aligned to the Human transcriptome and genome with TopHat. The genome version was GRCh38, with annotations from Ensembl release 99. Quantification was done with htseq-count. Normalization and differential expression analysis were done with the DESeq2 package. Genes with a sum of counts less than 10 over all samples were filtered out, then size factors and dispersion were calculated. Normalized counts were used for several quality control assays, such as counts distributions and principal component analysis, which were calculated and visualized in R. Pair-wise comparisons were tested with default parameters, except not using the independent filtering algorithm. Significance threshold was taken as padj<0.05 (default). In addition, significant genes were further filtered by the log2FoldChange value. This filtering was baseMean-dependent and required a baseMean above 5 and an absolute log2FoldChange higher than 5/sqrt(baseMean) + 0.6 (for highly expressed genes this means a requirement for a fold-change of at least 1.5, while genes with a very low expression would need a 7-fold change to pass the filtering). Finally, results were combined with gene details (such as symbol, known transcripts, etc.), taken from the results of a BioMart query (Ensembl, release 99), to produce the final Excel file.

#### Data Mining in Lung Cancer Cohort

Gene expression in lung cancer was taken from the combined cohort of TCGA, TARGET and GTEx. And was analyzed using the UCSC Xena browser (http://xena.ucsc.edu/). The genes co-expressed with levels of IGF2BP1 were separated to high and low IGF2BP1, and are presented in a violin plot.

#### Sample preparation for MS analysis

H1299 cells were treated with 1.5µM AVJ16/DMSO for 96 hours. Cell pellet was homogenized in RIPA buffer containing protease and phosphatase inhibitors, clarified by centrifugation, and the supernatant was subjected to protein precipitation by the chloroform/methanol method^34^. The precipitated proteins were solubilized in 100 μl of 8M urea, 10 mM DTT, 25 mM Tris-HCl pH 8.0 and incubated for 30 min at 22°C. Iodoacetamide (55 mM) was added followed by incubation for 30 min (22°C, in the dark), followed by re-addition of DTT (10 mM). 25 μg of protein was transferred into a new tube, diluted by the addition of 7 volumes of 25 mM Tris-HCl pH 8.0 and sequencing-grade modified Trypsin (Promega Corp., Madison, WI) was added (0.4 μg/ sample) followed by incubation overnight at 37°C with agitation. The samples were acidified by addition of 0.2% formic acid and desalted on C18 home-made Stage tips. Peptide concentration was determined by Absorbance at 280 nm and 0.75 µg of peptides were injected into the mass spectrometer.

#### nanoLC-MS/MS analysis

MS analysis was performed using a Q Exactive-HF mass spectrometer (Thermo Fisher Scientific, Waltham, MA USA) coupled on-line to a nanoflow UHPLC instrument, Ultimate 3000 Dionex (Thermo Fisher Scientific, Waltham, MA USA). Peptides dissolved in 0.1% formic acid were separated without a trap column over a 120 min acetonitrile gradient run at a flow rate of 0.3 μl/min on a reverse phase 25-cm-long C18 column (75 μm ID, 2 μm, 100Å, Thermo PepMapRSLC). The instrument settings were as described^35^. Survey scans (300–1,650 m/z, target value 3E6 charges, maximum ion injection time 20 ms) were acquired and followed by higher energy collisional dissociation (HCD) based fragmentation (normalized collision energy 27). A resolution of 60,000 was used for survey scans and up to 15 dynamically chosen most abundant precursor ions, with “peptide preferable” profiles, were fragmented (isolation window 1.8 m/z). The MS/MS scans were acquired at a resolution of 15,000 (target value 1E5, maximum ion injection times 25 ms). Dynamic exclusion was 20 sec. Data were acquired using Xcalibur software (Thermo Scientific). To avoid a carryover and to equilibrate the C18 column, the column was washed with 80% acetonitrile, 0.1% formic acid for 25 min between samples, as per standard protocol.

Raw data were processed using MaxQuant (MQ) version 1.6.5.0^36^, and the embedded Andromeda search engine^37^. The bioinformatics was performed with Perseus suite (version 1.6.2.3). The data were filtered for reverse, contaminants and identified by site. Then the data were filtered such that a protein had to have non-zero LFQ intensity in all 8 samples with 3 or more peptides. The significantly enriched proteins were found (two-sample t test with a permutation-based FDR method) and further selected using an adjusted p value.

#### Pathway enrichment analysis

Pathway enrichment analysis was used to identify the signaling pathways that are differentially regulated in treated vs nontreated populations. Changes in FDR<0.05 and Log_2_fold change >0.5 or <-0.5 were considered significant and analysed to see enrichment in pathways, using KEGG 2021 that was run using enrichR. Pathway enrichment analysis and visualization of omics data using GSEA and Cytoscape was done as described^38^, shortly: GSEA Analysis was performed for all human gene sets, the all human gene set collection was obtained from http://baderlab.org/GeneSets. Only significant pathways (FDR<0.05) were plotted into Cytoscape using “EnrichmentMap” plugin. Pathways were connected by lines if they shared many genes. Network layout and clustering algorithms automatically grouped redundant and similar pathways into major biological themes and collapsed into a single biological theme.

#### Statistics and reproducibility

At least three biological repeats were used for each experiment, and each experiment was performed at least twice. For each RT-PCR experiment, 3 biological repeats were performed, with 3 technical repeats for each sample. eCLIP was done in duplicate RNA-seq was done in triplicates and four biological repeats were analyzed by MS for each condition. Mice xenografts were performed three time with at least five male and five female mice for DMSO, and five male and five female mice injected with AVJ16. Data are shown as mean ± SEM unless otherwise stated.

## RESULTS

### AVJ16 binds specifically to IGF2BP1 in cells

Using both a microscale thermophoresis assay as well as NMR analysis, we demonstrated that AVJ16 binds to IGF2BP1 protein in-vitro^14^. Target engagement in cells, however, presents additional complexities, including membrane barriers and a crowded internal protein environment.^39^ To confirm direct binding of AVJ16 to IGF2BP1 in cells, a cell thermal shift stability assay (CETSA)^40^ was performed using the human lung cancer cell line H1299 that expresses high levels of IGF2BP1 endogenously. This assay is dependent on both uptake of the compound by cells and subsequent binding to its target protein in vivo. Compound binding to a protein generally enhances the thermal stability of the protein (Tm) that can be measured by quantifying the amount of soluble protein remaining after heating and lysing cells. Proteins are then electrophoresed on SDS-PAGE gels and visualized using target-specific antibodies. As seen in Fig. 1, AVJ16 interactions with IGF2BP1 stabilize the protein, resulting in an increase in its Tm by two degrees (from 46°C to 48°C) (Fig 1A, B). AVJ16 specificity for IGF2BP1 is also evident from the fact that no change in the Tm for IGF2BP2 and IGF2BP3 (47.5°C and 46°C, respectively) is observed (Fig 1A, C-D). These findings demonstrate not only that AVJ16 is capable of penetrating cells in culture but also specifically binds IGF2BP1, while exhibiting no significant interaction with the other paralogs.

**Figure 1.**
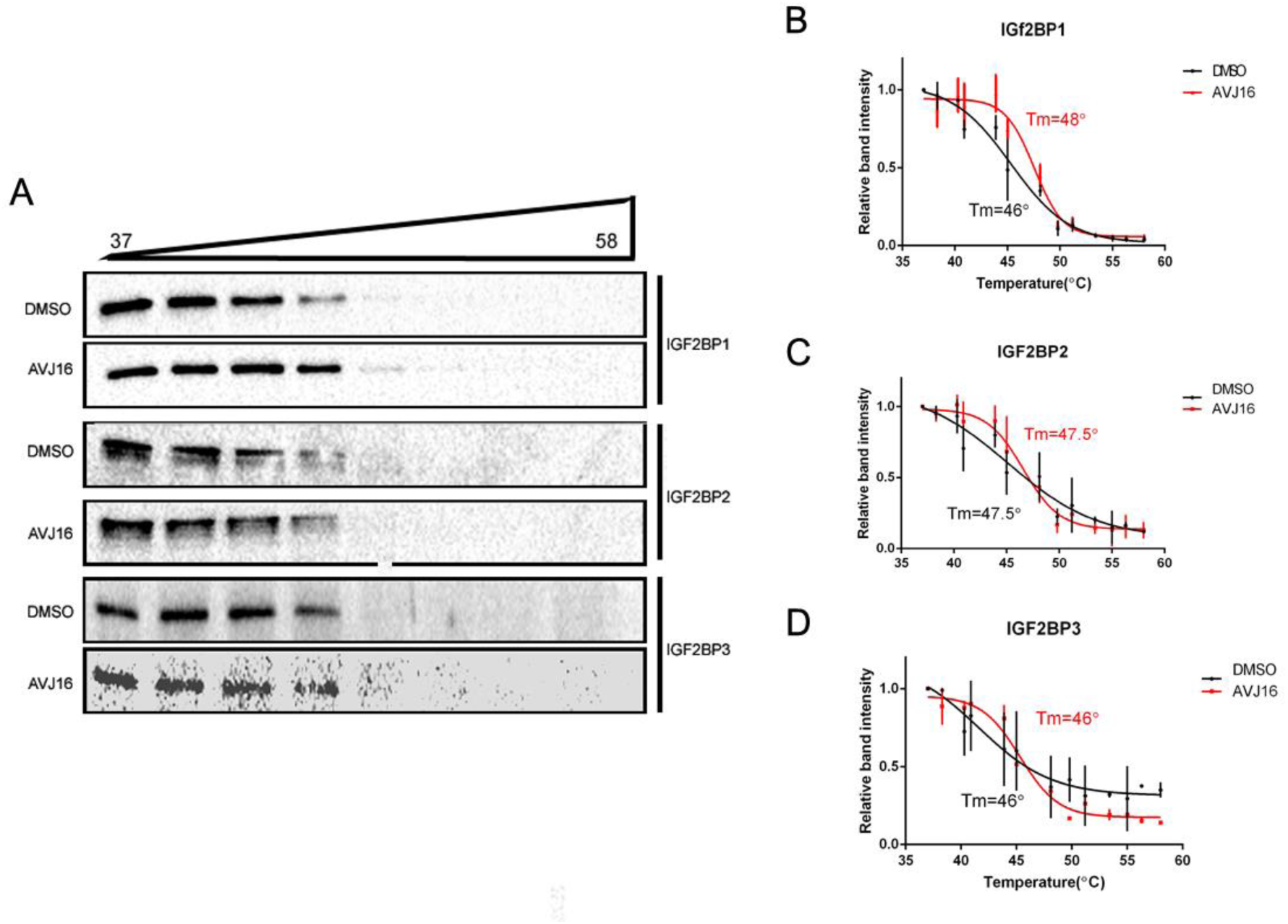
AVJ16 binds to IGF2BP1 in cells. CETSA was performed using H1299 cells. (A) Cells cultured for 3 hours with either 10μM AVJ16 or DMSO were subjected to the indicated temperature gradient for 3 minutes, lysed, and the soluble protein electrophoresed and analyzed on Western blots. Protein was visualized using antibodies specific for IGF2BP1, 2, or 3 (as indicated). The fraction of each IGF2BP paralog remaining intact was graphed as a function of temperature: (B) IGF2BP1; (C) IGF2BP2; (D) IGF2BP3. A Tm change of two degrees is observed in IGF2BP1, but no shift in Tm is observed in IGF2BP2 or 3. Three biological repeats were performed.

### AVJ16 directly perturbs RNA binding

Given that IGF2BP1 is a direct target of AVJ16, we reasoned that profiling the mRNAs whose binding to IGF2BP1 was affected by AVJ16 treatment could help in understanding the mechanism by which the inhibitor is effective against LUAD. To identify mRNAs bound directly to IGF2BP1, we performed, in conjunction with the company eCLIPSE Bioinnovation, an eCLIP (enhanced CrossLinking -ImmunoPrecipitation) analysis.^17^ Intact H1299 cells were UV irradiated to crosslink proteins and associated RNAs, and IGF2BP1 was pull downed via RNA Immunoprecipitation (RIP) with an anti-IGF2BP1 antibody. The associated RNAs were analyzed using deep sequencing, and the resulting peaks (reads) were mapped to their corresponding RNAs. Our analysis revealed that 22,311 peaks, originating from 5181 genes, were statistically significant targets of IGF2BP1. (A peak was defined as statistically significant if it had a log_2_ fold enrichment > 3 and p-value < 0.001). More than 70% of these targets were located in the 3’UTR, while only 20% were in the CDS, consistent with data from other eCLIP experiments with IGF2BP1 in different cell lines^17^ (Supplemental Data File 1; Figure 2A).

**Figure 2.**
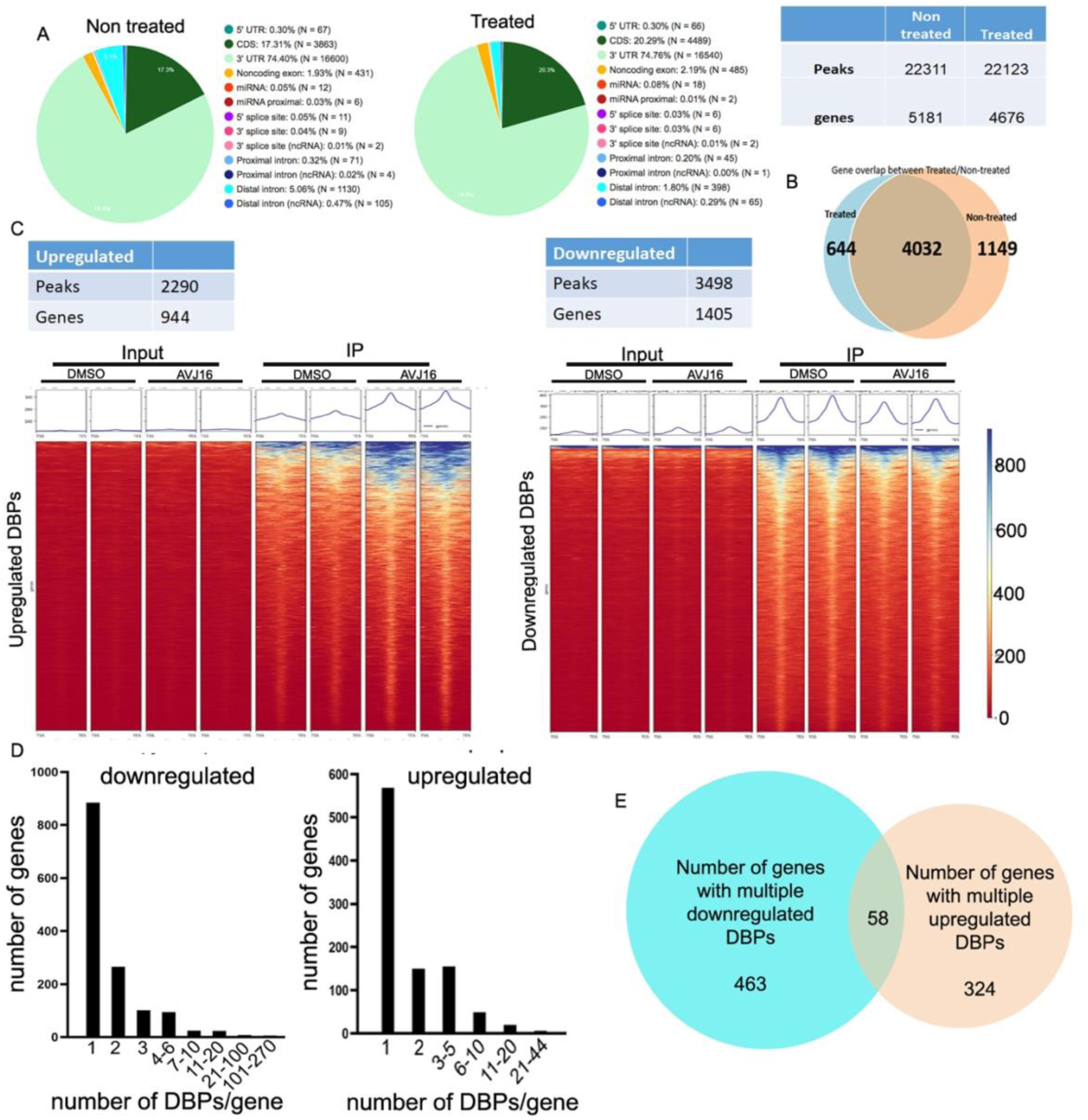
Evaluation of RNA targets of IGF2BP1 using eCLIP. eCLIP was performed using H1299 cells treated for 24 hours with either DMSO or 1.5µM AVJ16 in duplicate. (A) Pie charts showing the distribution of the peaks from DMSO (Non-treated) or AVJ16 (Treated) cells according to the region in the RNA to which they map. The table to the right shows the number of peaks and genes in which they are found. (B) Venn diagram showing the overlap of genes containing eCLIP peaks from DMSO and AVJ16-treated cells. (C) Tables showing the quantification of peaks and the genes in which they are found that were significantly up or down regulated as a result of AVJ16 treatment. The regions surrounding these differentially bound peaks (DBPs) were scaled to the same size (based on their transcription start site (TSS) and transcription end site (TES)) and calculated scores per genome region were plotted as a heat map, highlighting the peaks of the eCLIP compared to input, and comparing DMSO vs. AVJ16-treated peaks. (D) The number of genes (Y axis) with the indicated number of DBP peaks per gene (X axis) was plotted separately for the downregulated or upregulated DBPs. (E) In genes containing multiple DBPs, the large majority of genes (785/837=93.8%) are either all downregulated or all upregulated, as represented in the Venn diagram.

Upon treatment of H1299 cells with 1.5µM AVJ16, 22,123 statistically significant peaks originating from 4676 genes were identified. Remarkably, we observed a very substantial overlap of 4032 genes between AVJ16-treated and untreated samples, indicating the robustness and reproducibility of the eCLIP assay. Compared to the untreated samples, 1149 genes were no longer bound after treatment, while an unexpected 644 genes were newly bound upon treatment (Figure 2B)

The eCLIP data allows us not only to see which RNAs are directly interacting with IGF2BP1 but also to identify motifs enriched in IGF2BP1-bound RNAs. Despite the fact that these sequences are not necessarily the actual protein recognition site but only present in the crosslinked oligonucleotide, a comparison of enriched motifs identified by HOMER analysis reveals that the first, most common motif in both AVJ16 and DMSO-treated cells is the same (UUUCCGAA; Supplemental Fig. 1). Although there is some variation in the motifs observed less frequently in the eCLIP’d RNAs, certain similarities are present (e.g., UCCAG, CCXGUU, GXGGCCC, UCCUXXUGG). These results are perhaps not surprising in light of the high overlap of genes bound by IGF2BP1 in both AVJ16 and DMSO-treated cells.

We sought to assess the differential binding of IGF2BP1 to RNA peaks. A peak was considered a differentially bound peak (DBP) if there was a statistically significant difference in read counts between treated and untreated conditions (see Materials and Methods). Out of all peaks, 3498 were downregulated and 2290 upregulated, originating from 1405 and 944 genes, respectively. When all DBPs are scaled to the same size, and calculated scores per genome region are plotted as a heat map highlighting the DBPs, global down and up regulation of DBPs compared to the DMSO control can be observed. These results demonstrate that IGF2BP1 RNA interactions are regulated upon AVJ16 treatment (Supplemental Data File 1; Figure 2C).

Out of all the genes with down regulated DBPs, 884 contained a single DBP, and 521 genes contained multiple DBPs. Of the genes containing upregulated DBPs, 562 had a single DBP, and 382 genes had multiple DBPs (Figure 2D). Understanding how the binding of IGF2BP1 to specific sites influences gene expression in genes with multiple peaks presents a challenge, as we do not know whether multiple binding events have a combinatorial effect, or which peaks have a functional or stabilizing effect. To address this issue, we assessed the degree of overlap between genes containing either downregulated or upregulated peaks. Strikingly, we found that out of the 2349 genes with DBPs, only 58 genes contained both up and down-regulated DBPs, which represents only 2.5% of the genes (Fig 2E). In other words, DBPs within a given gene tend to behave in the same way, being either up or down-regulated together by the treatment. This finding suggests that, to a first approximation, the overall effect of IGF2BP1 binding to genes can be considered binary, without having to assess each individual peak. We can then conclude that 1) the predominant effect of AVJ16 treatment on H1299 cells is perturbation of the direct binding of IGF2BP1 to many target RNAs, but 2) there are also many other RNAs that become directly bound to IGF2BP1 as a result of the treatment.

### Large transcription changes upon IGF2BP1 inhibition

To investigate the impact of AVJ16 treatment on the steady state of the RNAs in H1299 cells, RNA-seq was conducted after treating with a low dose (1.5µM) of AVJ16 or DMSO for 48 hours. Differentially expressed genes (DEGs) were statistically identified (Supplemental Data File 1), and principal component analysis (PCA) revealed that the treated and DMSO-treated cells segregated into two different compartments (PC1 54%, PC2 32%) (Supplemental Figure 2A). Among the significantly affected genes, 1306 were downregulated (log_2_ fold change < -0.5, p_adj_ < 0.1) and 823 were upregulated (log_2_ fold change > 0.5, p_adj_ < 0.1). The most significantly affected genes are shown in Figure 3A.

**Figure 3.**
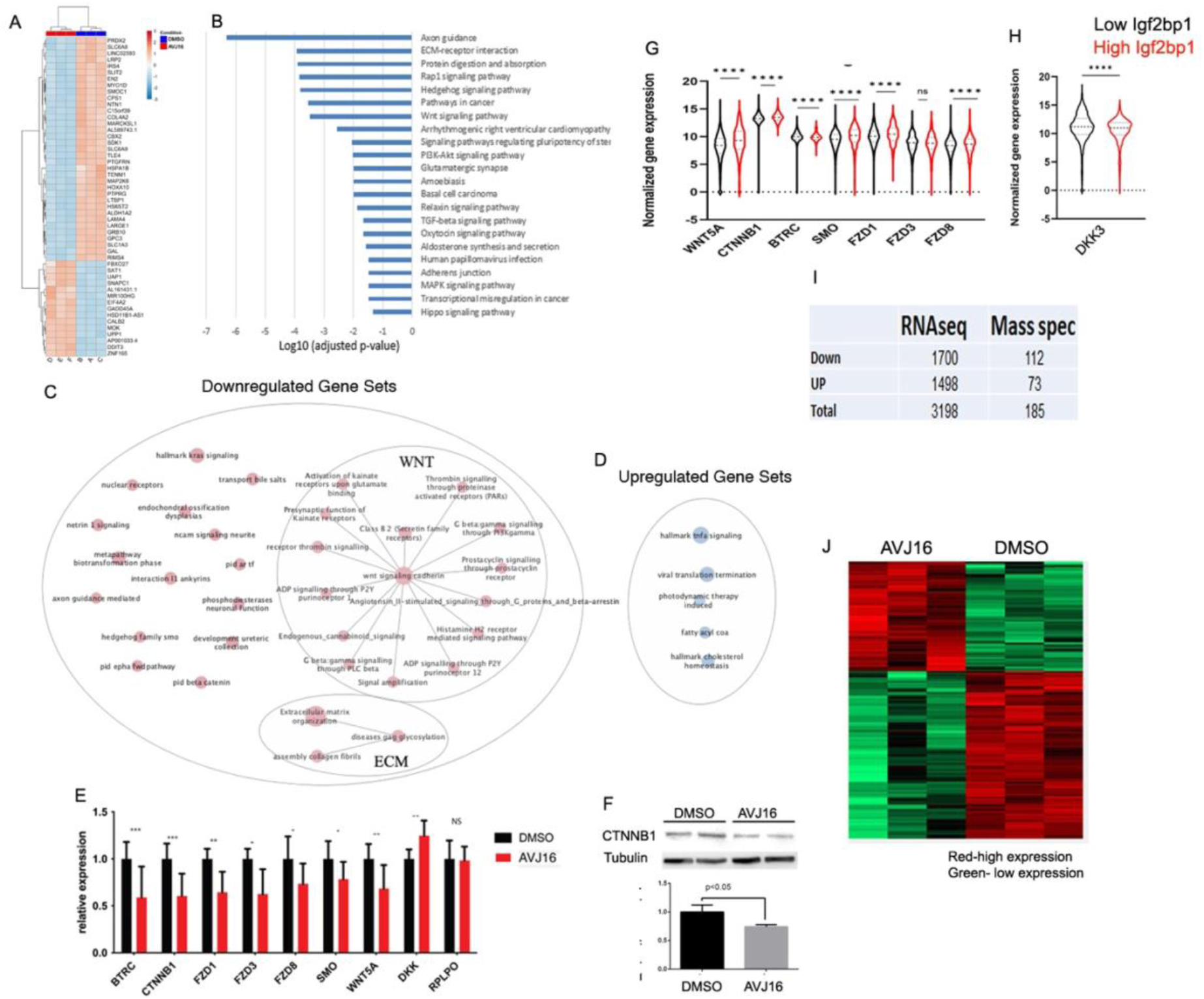
Impact of AVJ16 treatment on the steady state of RNAs in H1299 cells. RNA-seq was performed on RNA from cells treated with 1.5µM AVJ16 or DMSO for 48 hours. (A) Heat map of the most significantly affected genes after AVJ16 treatment, with downregulated genes shown in blue and upregulated genes shown in red. (B) KEGG annotation of gene sets downregulated by AVJ16 treatment, showing significant enrichment in IGF2BP1-related pathways, such as axon guidance, ECM-receptor interactions, WNT signaling, and PI3K-Akt signaling. (C) Enrichment map of the AVJ16-downregulated genes analyzed by Gene Set Enrichment Analysis (GSEA) for all human gene sets, showing significant enrichment of pathways associated with transformed phenotypes and signaling pathways activated during cancer progression, with WNT and ECM being the most clustered pathways. (D) Enrichment map of the AVJ16-upregulated genes analyzed by GSEA for all human gene sets. (E) RNA from cells treated with 1.5µM AVJ16 or DMSO for 48 hours was analysed for expression of WNT-related genes by qPCR. The seven WNT-related genes downregulated in RNAseq were significantly downregulated in H1299 cells upon AVJ16 treatment, compared to the DMSO-treated control group (RPLPO was added as a housekeeping gene control). The WNT inhibitor, DKK3, was upregulated in the RNAseq analysis, and its expression was significantly upregulated in the AVJ16-treated H1299 cells. (F) Western blot analysis of beta catenin protein levels in H1299 cells treated with 1.5µM AVJ16 or DMSO for 48 hours. Beta catenin protein levels were downregulated, consistent with the reduction in its RNA level observed in the RNAseq data. (G-H) Correlation between the expression of WNT pathway genes and IGF2BP1 expression was analyzed in data collected from human databases. Gene expression data came from the combination of TCGA, TARGET and GTEx lung cohorts and show a significant correlation for all the genes examined, with the exception of FZD3. (G) Genes promoting WNT signaling are upregulated when IGF2BP1 is high. (H) DKK3A RNA is downregulated when IGF2BP1 is high. (I) Table comparing changes in RNA levels (RNAseq) and in protein levels (Mass spec) in H1299 cells treated with 1.5µM AVJ16 or DMSO for either 48 hours (RNAseq) or 96 hours (Mass spectrometry). (J) Heat map of all statistically significant upregulated and downregulated proteins.

Downregulated genes included many known targets of IGF2BP1, such as βTrCP1, GLI1, and ABC transporters (shown in a volcano plot representation in Supplemental Figure 2B). KEGG annotation of gene sets downregulated by AVJ16 revealed that many well-known IGF2BP1-related pathways were significantly enriched, such as axon guidance, ECM-receptor interactions, and WNT and PI3K-Akt signaling (Figure 3B). Strikingly, although over a third of the DEGs were upregulated, no pathways were statistically enriched in these genes, suggesting that the upregulated genes may constitute more of a random collection rather than a coordinated set of pathways. To ensure that AVJ16 does not upregulate cancer-relevant pathways, Gene Set Enrichment Analysis (GSEA) was performed, using the all-human gene set collection obtained from http://baderlab.org/GeneSets. In the representation shown in Figure 3C, pathways are depicted as nodes (red nodes, downregulated gene sets, and blue nodes, upregulated gene sets), connected by lines if the pathways share many genes. Network layout and clustering algorithms automatically grouped redundant and similar pathways into major biological themes, and collapsed them into a single biological theme. The analysis indicates that many RNAs downregulated by AVJ16 treatment are significantly enriched in gene sets associated with transformed phenotypes and signaling pathways activated during cancer progression, in accordance with known roles of IGF2BP1 in lung carcinoma. Many of the pathways from the KEGG annotation are present in the GSEA, but WNT and ECM are the two most clustered pathways (Figure 3C). Although the GSEA analysis did identify a small number of significantly enriched gene sets among the upregulated genes, none of these are associated with a direct effect on tumorigenesis (Figure 3D).

Previous studies have demonstrated that IGF2BP1 enhances WNT pathway signaling and that downregulation of IGF2BP1 can reduce the resistance of chemotherapy-resistant cancer cells that have an activated Wnt/β-catenin signaling pathway in colorectal carcinomas (CRC).^41,42^ In order to confirm the RNA-seq results and assess the significance of IGF2BP1 in the WNT pathway in LUAD, we performed qPCR to quantify the expression levels of seven genes involved in the WNT pathway that were down regulated in the RNA-seq (Supplemental Figure 2B). Expression of these WNT-related genes is reduced by incubation of H1299 cells with AVJ16 (Figure 3E), but no reduction in these RNAs is observed in AVJ16-treated RKO cells, which do not express IGF2BP1 (Supplemental Figure 2C). Notably, DKK3, a WNT inhibitor, is upregulated in H1299 treated cells, leading to further inhibition of the WNT pathway. As expected, expression of β-catenin protein is downregulated concomitant with its RNA levels (Figure 3F).

Using the data from the human TCGA TARGET GTEx lung data bases, we explored the potential correlation between IGF2BP1 expression and genes associated with the WNT pathway. All of the WNT pathway genes as defined by KEGG annotation and that were differentially expressed in the RNA-seq analysis were significantly associated with IGF2BP1 expression, with the exception of one gene, FZD3, which showed no significant change in expression (Figure 3G, H). This motivated us to examine global proteome changes upon AVJ16 treatment. H1299 cells were treated with 1.5µM of AVJ16 for 96 hours, and the cell lysates were analysed using mass spectrometry (MS). Although the total number of identifiable proteins from MS studies is typically small compared to the identified RNAs in comparable RNAseq experiments, our analysis revealed that 112 proteins were significantly downregulated and 73 were significantly upregulated (Supplemental Data File 1; Figure 3I, J), indicating that a global change was induced at the proteome level upon treatment.

### Correlation between eCLIP, RNA-seq, and MS

RNA binding proteins can affect the bound RNAs via numerous mechanisms, such as RNA stability, translation, splicing, and intracellular localization. If the major effect of AVJ16 treatment is mediated by affecting binding of IGF2BP1 to its target RNAs, we reasoned that significant changes in RNA binding caused by AVJ16 treatment would be reflected in altered steady state RNA and protein levels. Comparison of the log_2_ fold changes of the RNAseq data and the eCLIP data on a gene-by-gene basis reveals a strong, positive correlation (Mann-Whitney unpaired t-test p<0.0001, χ_2_<0.0001) (Figure 4A), suggesting that the effect of AVJ16 on IGF2BP1’s binding to a given RNA is generally reflected in a comparable change in its steady state RNA level, and ultimately in expression of its encoded protein.

**Figure 4.**
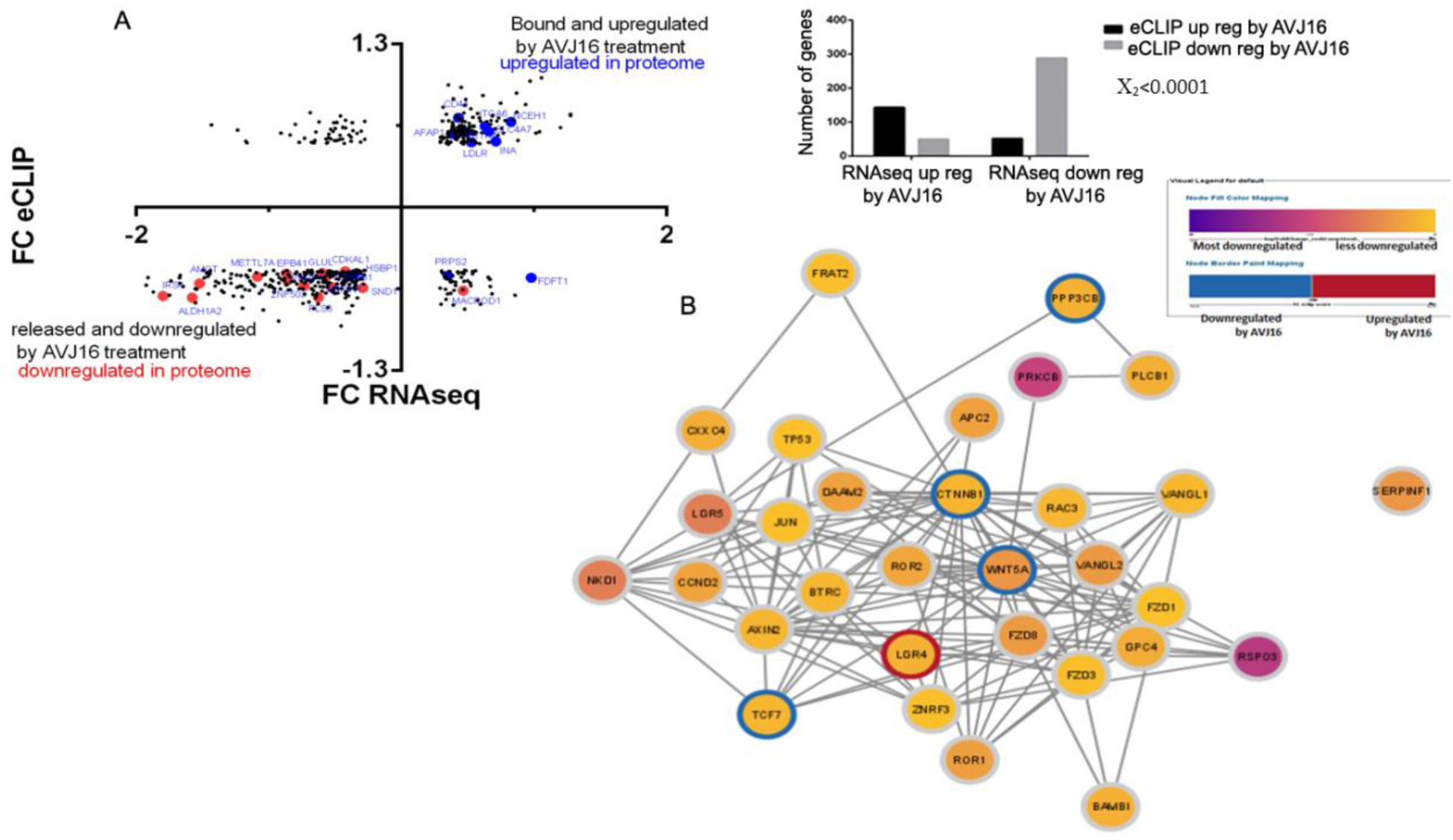
Correlation between eCLIP, RNAseq, and MS data. (A)The log_2_ fold changes of the RNAseq data (Y axis) and the log_2_ fold changes of the eCLIP data (X axis) were compared on a gene-by-gene basis (black dots). Strong, positive correlation is indicated (Mann-Whitney unpaired t-test p<0.0001, χ_2_<0.0001). The graph to the right quantifies the distribution of the genes up or down regulated in RNAseq and their up or down regulation as analyzed by eCLIP. Overlaid on the distribution plot (left) are those genes upregulated in the MS data (blue dots) and those downregulated in the MS data (red dots). (B) Mapping of downregulated WNT pathway DEGs based on RNAseq data onto a protein-protein interaction (PPI) network. The circle surrounding each node indicates whether the direct binding to the gene was up (red) or downregulated (blue) by AVJ16 treatment. The color of the nodes represents the degree of downregulation in RNA levels. Genes whose direct binding to IGF2BP1 is downregulated by AVJ16 (WNT5A, CTNNB1, PPP3CB, TCF7) and LGR4, whose binding is upregulated, play a major role in influencing the WNT PPI network.

To analyze this behavior in greater detail, we mapped the downregulated WNT pathway DEGs from the RNAseq data, onto a protein-protein interaction (PPI) network. The colour of the circle surrounding the node indicates whether the direct binding to the gene was up- or down-regulated by AVJ16 treatment, thus graphically distinguishing between direct targets of IGF2BP1 and the secondary effect on the downstream transcriptome. As indicated in the WNT network, there are four genes whose direct binding to IGF2BP1 is downregulated by AVJ16 (WNT5A, CTNNB1, PPP3CB, TCF7), and one (LGR4) that is upregulated. These genes influence the WNT PPI network and lead to lower RNA expression and protein levels of other members in the pathway (Figure 4B).

In conclusion, these experiments provide direct evidence that AVJ16 induces significant perturbations in RNA binding, as observed in the eCLIP experiments, resulting in substantial changes at both the transcriptome and proteome levels. Pathway analysis reveals that AVJ16 treatment leads to downregulation of IGF2BP1-related pathways such as the WNT pathway, a pathway implicated in lung cancer tumorigenesis.

### AVJ16 inhibits tumorigenic properties of cancer cells in an IGF2BP1-dependent manner

The human lung cancer cell line H1299 expresses high levels of IGF2BP1. We previously observed that AVJ16 is very effective at inhibiting cell proliferation in these cells^14^. Colony formation and anchorage-independent growth are considered to be hallmarks of cancer and neoplastic cells.^27^ To investigate the effect of AVJ16 on these processes, cells were cultured with or without 4µM AVJ16 at low density to allow colony formation from single cells. As seen in Figure 5A, AVJ16 reduced colony formation in H1299 cells by over 40%. To assess the specificity of this effect, we made use of a mouse lung carcinoma line, LKR-M, that has been selected to be highly metastatic but does not express endogenous IGF2BP1^4,28^. LKR-M cells stably transfected with full length IGF2BP1 (LKR-M-Fl) are similarly inhibited in colony formation when treated with AVJ16 (Figure 5B), but this treatment has no effect on LKR-M cells lacking IGF2BP1 expression (LKR-M-GFP; Figure 5C). Colony formation in soft agar was significantly inhibited as well by AVJ16 in H1299 and LKR-M-Fl cells but not affected in LKR-M-GFP cells (Figure 5D-F).

**Figure 5.**
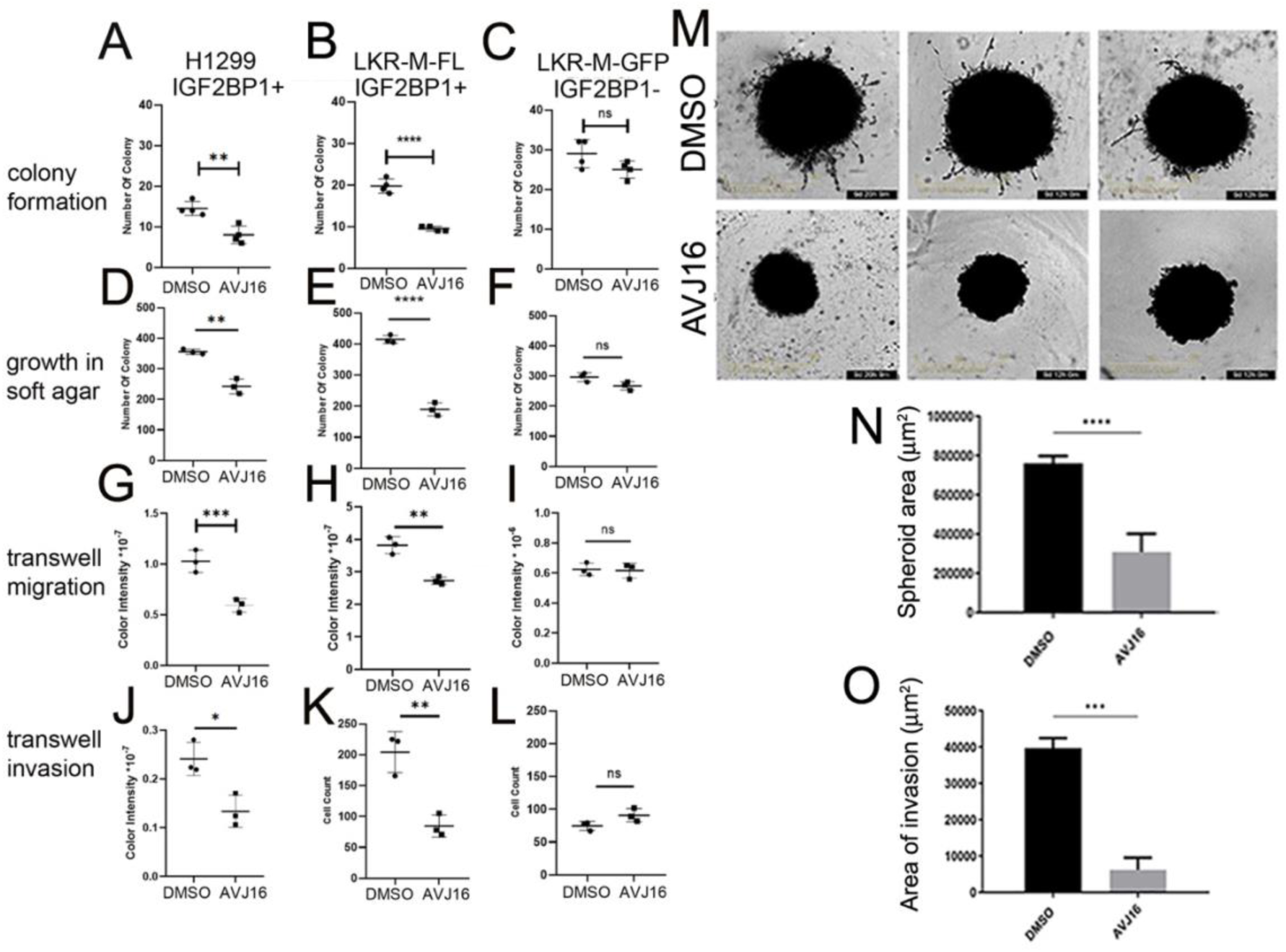
Effects of AVJ16 treatment on H1299 lung carcinoma cells. The effects of AVJ16 treatment (4μM) on colony formation (A,B,C), growth in soft agar (D,E,F), transwell migration (G,H,I) and transwell invasion (J,K,L), were compared to DMSO treatment in H1299 cells (A,D,G,J), LKR-M-Fl cells (B,E,H,K) and LKR-M-GFP cells (C, F, I, L). H1299 cells grown as spheroids embedded in Matrigel in the presence of 4μM AVJ16 or DMSO (M) were analyzed for growth (N) and invasion capacity (O). All experiments were done in triplicate.

Knockdown of IGF2BP1 has been shown to inhibit cell migration and invasion as well as tumor metastasis^4,21,43^. AVJ16 treatment of H1299 cells significantly reduces their migration in a transwell membrane assay (Figure 5G). Similarly, LKR-M-Fl cell migration is repressed upon AVJ16 treatment, whereas no significant reduction is observed in LKR-M-GFP cells (Figure 5H,I). Coating the membranes with Matrigel allows us to assess the ability of these cells to invade an environment similar to the extracellular matrix (ECM) that metastatic cells encounter in vivo. Here, too, AVJ16 significantly reduces invasion of cells expressing IGF2BP1 but not of those cells lacking IGF2BP1 (Figure 5J-L).

As 3D tumor cell culture is considered to more accurately reflect growth in the tumor microenvironment than 2D culture systems, especially when coupled with growth in an ECM-like solution^44^, we further tested the effect of AVJ16 on growth and invasion of H1299 cells as spheroids. AVJ16 significantly impedes both growth and invasion of H1299 cells in a 3D setting, particularly inhibiting the penetration of cells into the surrounding Matrigel (Figure 5M-O). In the absence of Matrigel, not only are the spheroids growth-inhibited by AVJ16 but they also demonstrate clearly visible cell death in a dose-dependent manner (Supplemental Figure 3A-C). These findings collectively support AVJ16’s potential as an effective inhibitor of tumor growth and invasion.

To assess AVJ16’s cytotoxicity, a lactate dehydrogenase (LDH) activity assay was performed. LDH is an enzyme released into the media after cellular damage or hemolysis and has been widely used to evaluate toxicity in cells and tissues. The LDH activity assay revealed that AVJ16 is not generally toxic to cells, even at high dosages (12 µM) and long incubation periods (96 hours; Supplemental Figure 3D).

### AVJ16 induces apoptosis

Numerous studies have highlighted the role of IGF2BP1 as an inhibitor of apoptosis (e.g.^10,41,45,46^). The appearance of cell death in the spheroids grown in Matrigel (Supplemental Fig. 3) suggest a potential induction of apoptosis upon AVJ16 treatment. To confirm this hypothesis, we employed a cleaved caspase 3/7 dye assay to detect caspase activity. The caspase activity in H1299 cells is significantly higher in the AVJ16-treated group compared to the non-treated group (Fig 6A), indicating that AVJ16 indeed induces apoptosis. It is important to note that caspase activity has been normalized here to cell number, inasmuch as AVJ16 also inhibited proliferation. Thus, although there are fewer cells in the treated group, the caspase activity per cell is higher. In addition, untreated H1299 cells have been noted to have a significant background caspase activity^47^, but nevertheless, AVJ16 dramatically increases this. Similar results were observed in LKR-M-Fl (Figure 6B), whereas low caspase activity was observed in LKR-M-GFP with or without AVJ16 treatment (Figure 6C). Furthermore, FACS analysis revealed that apoptosis is progressive and specific for IGF2BP1-expressing cells. At 24 hours, apoptosis is enhanced by AVJ16 almost 2.5-fold in H1299 cells (Supplemental Figure 4A), reaching a nearly 4-fold increase by 48 hours (Figure 6D-F). Increased γ-H2AX levels, indicative of double strand DNA breaks that are generated during apoptosis, further support this finding (Supplemental Figure 4B). FACS analysis also demonstrates that AVJ16 induction of apoptosis occurs in LKR-M-Fl cells but not in LKR-M-GFP cells (Figure 6G-L). Thus, the progressive and specific nature of apoptosis suggests that AVJ16 specifically induces pathways leading to cell death as opposed to general toxicity.

**Figure 6.**
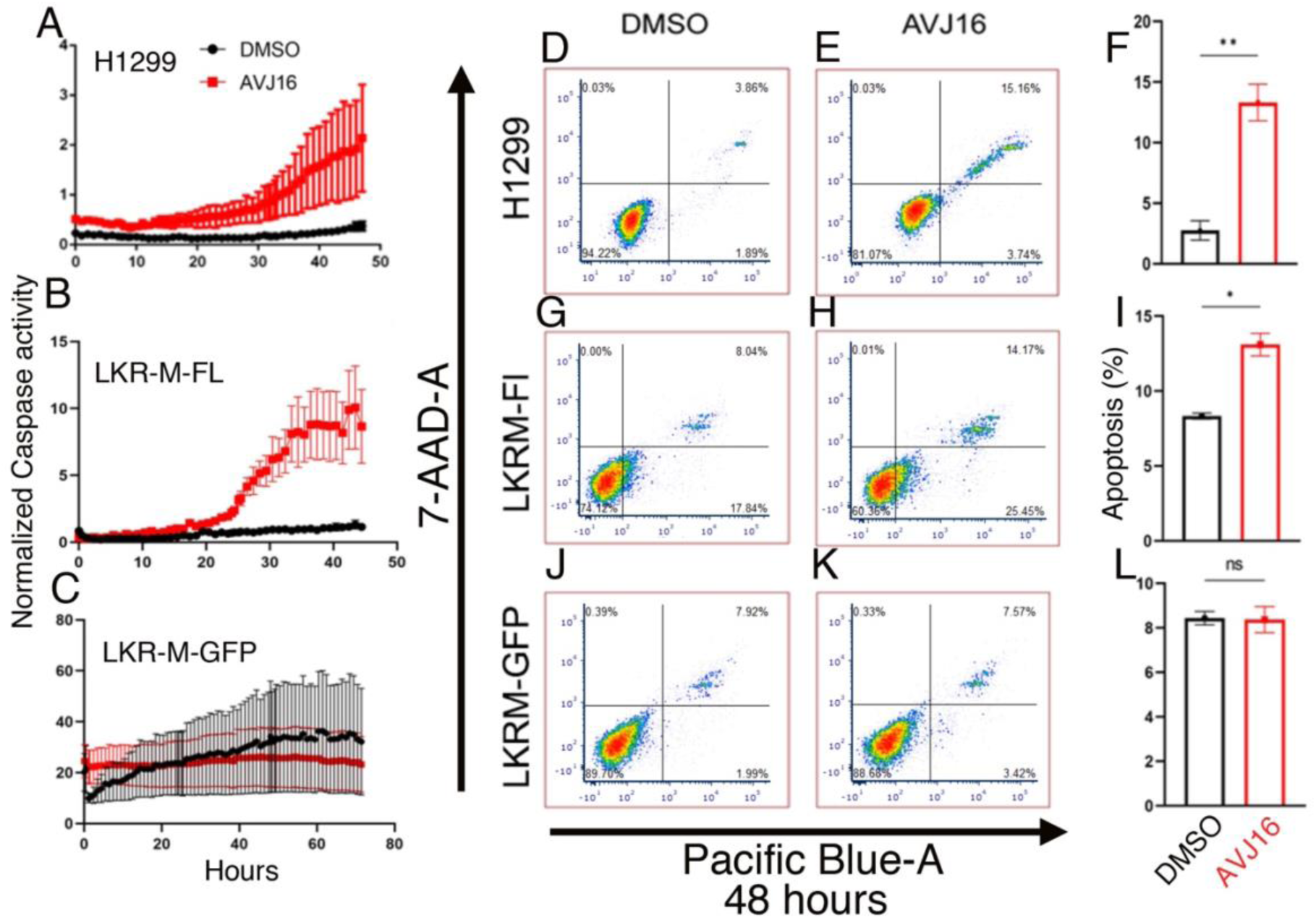
AVJ16 induces apoptosis. (A-C) Caspase 3/7 dye was added to cells treated with DMSO (black dots) or 4μM AVJ16 (red dots) and monitored by the IncuCyte® S3 Live-Cell Analysis System. The amount of detected dye was normalized to cell number and plotted as a function of time. (A) H1299. (B) LKR-M-FL cells. (C) LKR-M-GFP. (D,E,G,H,J,K) FACS sorting to was used to detect Annexin/7-AAD-A staining of cells treated with DMSO or 4μM AVJ16 for 48 hours. Late apoptotic cells appear in the top right quadrant. The percent of late apoptotic cells in DMSO vs. AVJ16-treated cells was compared for H1299 cells (F), LKR-M-FL cells (I), and LKR-M-GFP cells (L).

Collectively, the results obtained from these cell-based experiments demonstrate that AVJ16 binds to IGF2BP1 within cells, impedes growth, inhibits migration, and induces apoptosis and cell death specifically in cells that express IGF2BP1.

### AVJ16 treatment inhibits dye efflux from IGF2BP1-expressing cells

IGF2BP1 has been correlated with chemoresistance in different types of cancers, due to its ability to stabilize many pro-oncogenic RNAs, including ATP-binding cassette (ABC) transporters that are responsible for efficiently pumping drugs out of the cell.^45,48–50^ Dye efflux has been used as a readout for ABC transporter activity.^51,52^ By FACS analysis, we tested the ability of cells to efflux the nucleophilic dye, Hoechst 33342. A small side population is present in H1299 cells that resists dye accumulation (Fig. 7). AVJ16 treatment severely inhibits the ability of these cells to efflux the dye. A similar, although much smaller, side population of dye effluxing cells is observed in the LKR-M-FL cells, and AVJ16 treatment also causes this population to disappear. Addition of reserpine, an ABC transport blocker, produced a similar effect by abolishing the side population in both H1299 and LKR-M-FL cells. No such side population is evident in the control LKR-M-GFP cells that do not express IGF2BP1. These results suggest that cells expressing IGF2BP1 may have an enhanced ability to remove dye and presumably other chemicals taken up by the cell. Furthermore, AVJ16 can inhibit this ability, in an IGF2BP1-specific manner, potentially adding to its anti-tumorigenic repertoire.

**Figure 7.**
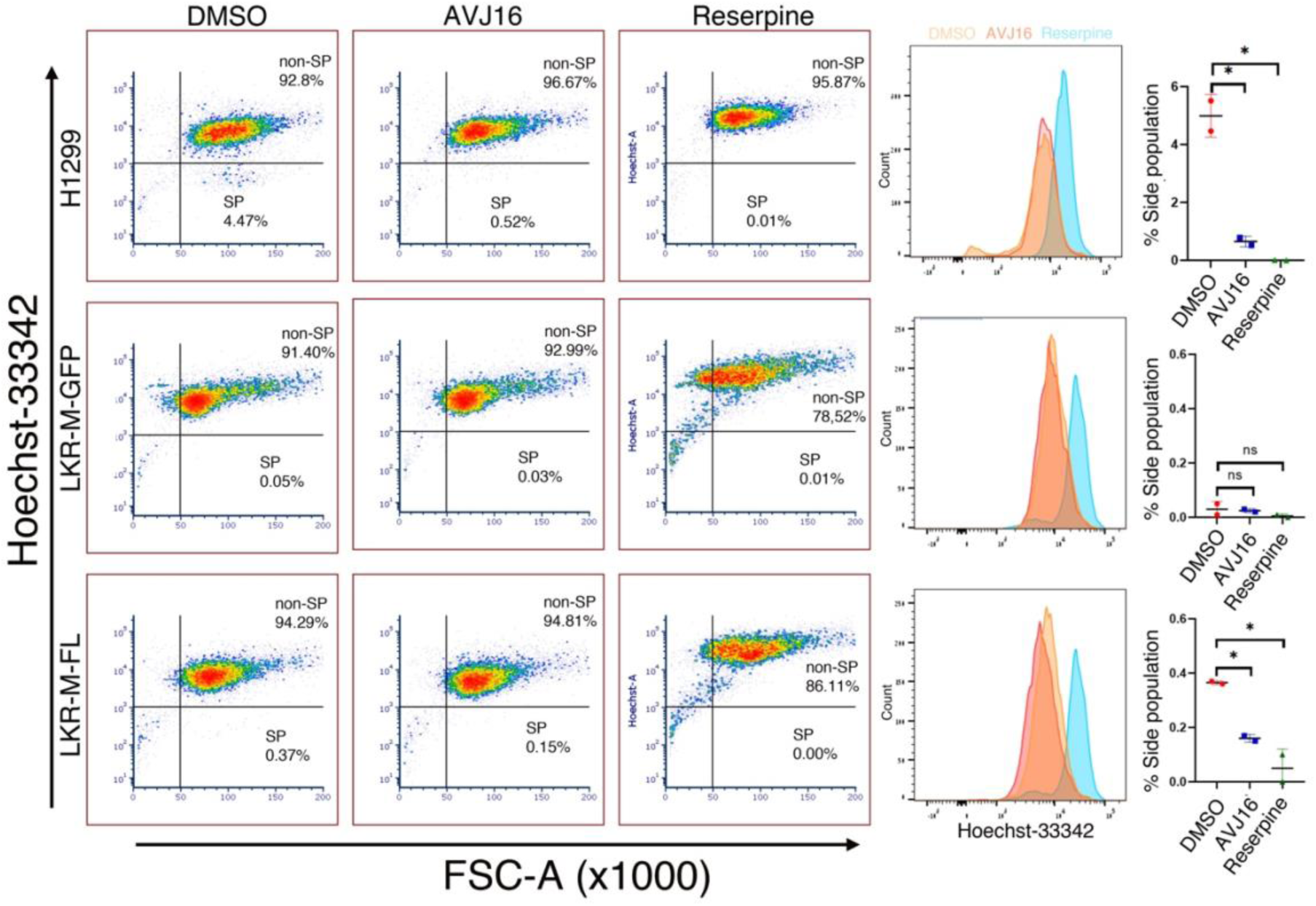
AVJ16 reduces dye efflux from H1299 cells. H1299, LKR-M-Fl, and LKR-M-GFP cells (as indicated on the left) were grown in the presence of DMSO, 4μM AVJ16, or 50μM Reserpine (as detailed in Materials and Methods), suspended in medium, incubated for 90 minutes with Hoechst-33342, and then sorted by FACS. The graphs on the right show the results from 2 biological repeats. The side population (*SP*) demonstrating dye efflux is seen in the bottom right quadrant.

### Syngeneic xenografts of IGF2BP1-expressing cells are inhibited by AVJ16

The efficacy of AVJ16 in the cell-based assays encouraged us to investigate the effectiveness of AVJ16 in vivo, using a mouse model. Previously^4^, we demonstrated that genetic inhibition of IGF2BP1 suppresses tumor progression and metastasis of LUAD. To evaluate the in vivo functions of AVJ16, we subcutaneously implanted LKR-M-Fl or LKR-M-GFP cells into syngeneic mice (Figure 8A). In the absence of AVJ16 treatment, LKR-M-Fl cells show a two to three-fold increase in tumor size compared to LKR-M-GFP, indicating that exogenous IGF2BP1 promotes tumor progression in these xenografts (compare Figure 8B,C with Supplemental Figure 5C). The maximum tolerated dose (MTD) of AVJ16 administered intraperitoneally (IP) was determined by our previously reported method^53,54^ to be 400 mg/kg body weight. When tumors became visible 12 days post-implantation, mice were injected IP with 25% of MTD (100 mg/kg), three times a week for three weeks. At around 30 days, mice were sacrificed (Figure 8A). No toxicity was observed, and there was no substantial change in mouse weight (Supplemental Figure 5A,B). Primary tumor growth of the LKR-M-Fl xenografts was almost completely inhibited in the AVJ16-treated mice (Figure 8B,C), while no effect on proliferation of the LKR-M-GFP xenografts was observed (Supplemental Figure 5C). In parallel experiments, mice were injected peritumorally (at a dose ∼500 times less than used for IP injections). Here as well, a significant reduction in primary tumor growth was observed (Supplemental Figure 5D,E), concomitant with fewer metastases to the lungs (Supplemental Figure 5F). To further determine the inhibitory effect of AVJ16, primary tumors were fixed in paraformaldehyde and stained for the expression of the cell proliferation marker, Ki67. Tumors from mice injected with AVJ16 showed a 40% reduction in nuclear Ki67 staining, as compared to those from DMSO-injected mice (Figure 8D). AVJ16, therefore, is an effective inhibitor of tumor growth and metastasis in mice.

**Figure 8.**
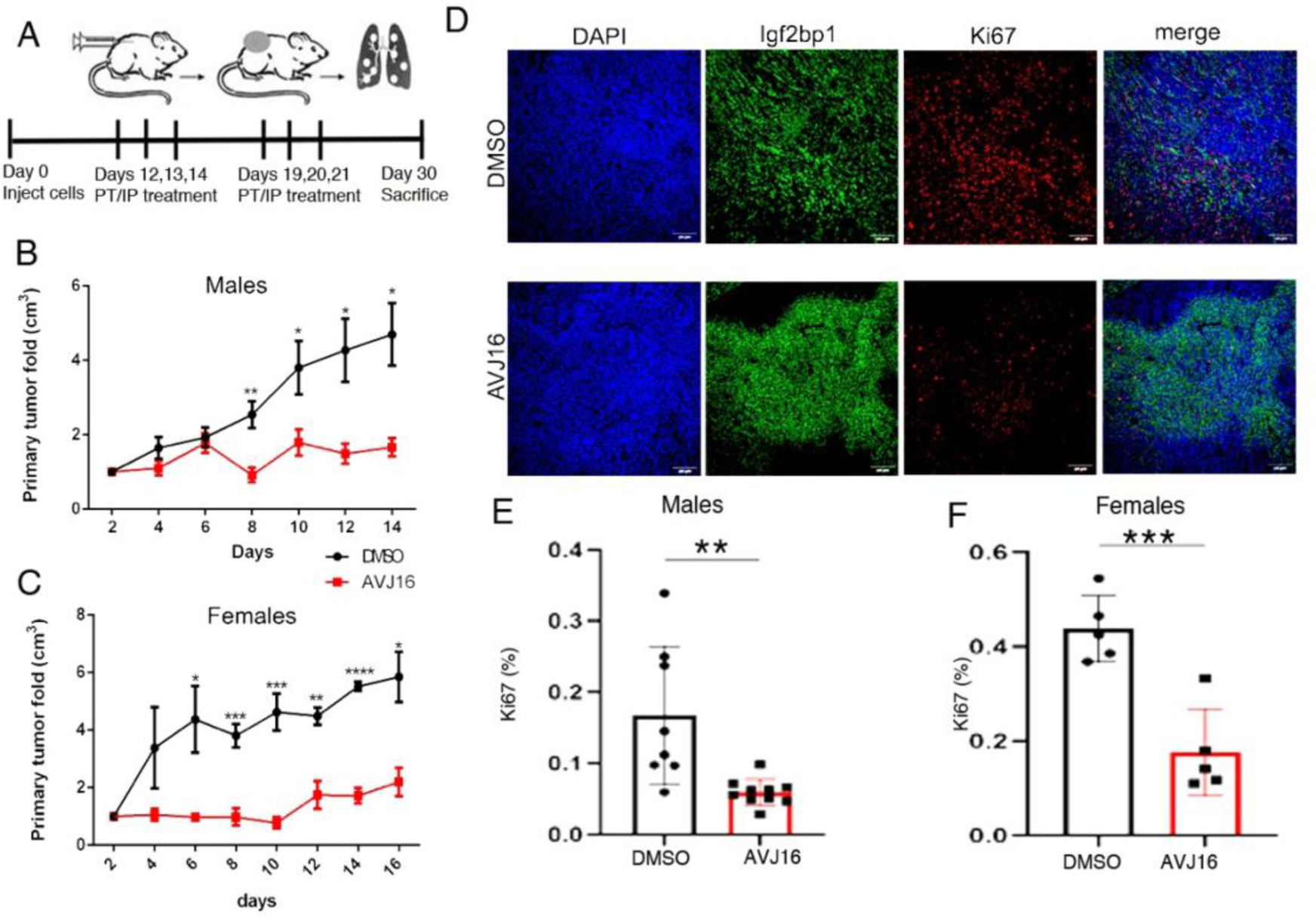
AVJ16 effect on mice xenograft model. (A) A schematic drawing of the injection protocol. Syngeneic mice were subcutaneously implanted with LKR-M-Fl cells and allowed to grow for 12 days, until the site of implantation was visible as a bulge. AVJ16 was injected intraperitoneally (IP, this figure) or peritumorally (PT, Supplemental Figure 4) at the times indicated below the timeline. The primary tumor fold increase was measured every two days (time 0, 12 days after subcutaneous injection of cells). Tumor growth was inhibited in both males (B) and females (C). The number of mice for all experiments was N≥5. (D) Representative images of primary tumors stained for DAPI, IGF2BP1 and Ki67. Ki67 positive nuclei were counted and normalised to the DAPI staining, showing more than 40% reduction of proliferation both in males (E) and in females (F). (N=3)

### Patient-derived tumor organoids from non-small cell lung carcinomas (NSCLC) are sensitive to AVJ16 treatment

Patient-derived organoids are three-dimensional structures formed from healthy or cancer cells isolated from a patient’s tissue, which self-organize to mimic the architecture and cellular interactions of the original tissue. Patient-derived tumor organoids retain the heterogeneity and specific characteristics of the patient’s cancer, making them valuable models for studying tumor behaviour and testing personalized treatments^55^. As proof of concept that AVJ16 could be a useful tool in a clinically relevant setting, we compared the response to AVJ16 treatment of a human healthy lung organoid versus a human NSCLC organoid from a surgically resected tumor that expresses IGF2BP1. The healthy lung organoids showed no elevated cell death when treated with either AVJ16 or DMSO (Figure 9A). The NSCLC-derived organoids expressing IGF2BP1, however, were strikingly sensitive to AVJ16 treatment, with elevated cell death observed at a concentration of 4μM (Figure 9B). These results underscore the specificity of AVJ16 for human lung tumors expressing IGF2BP1.

**Figure 9.**
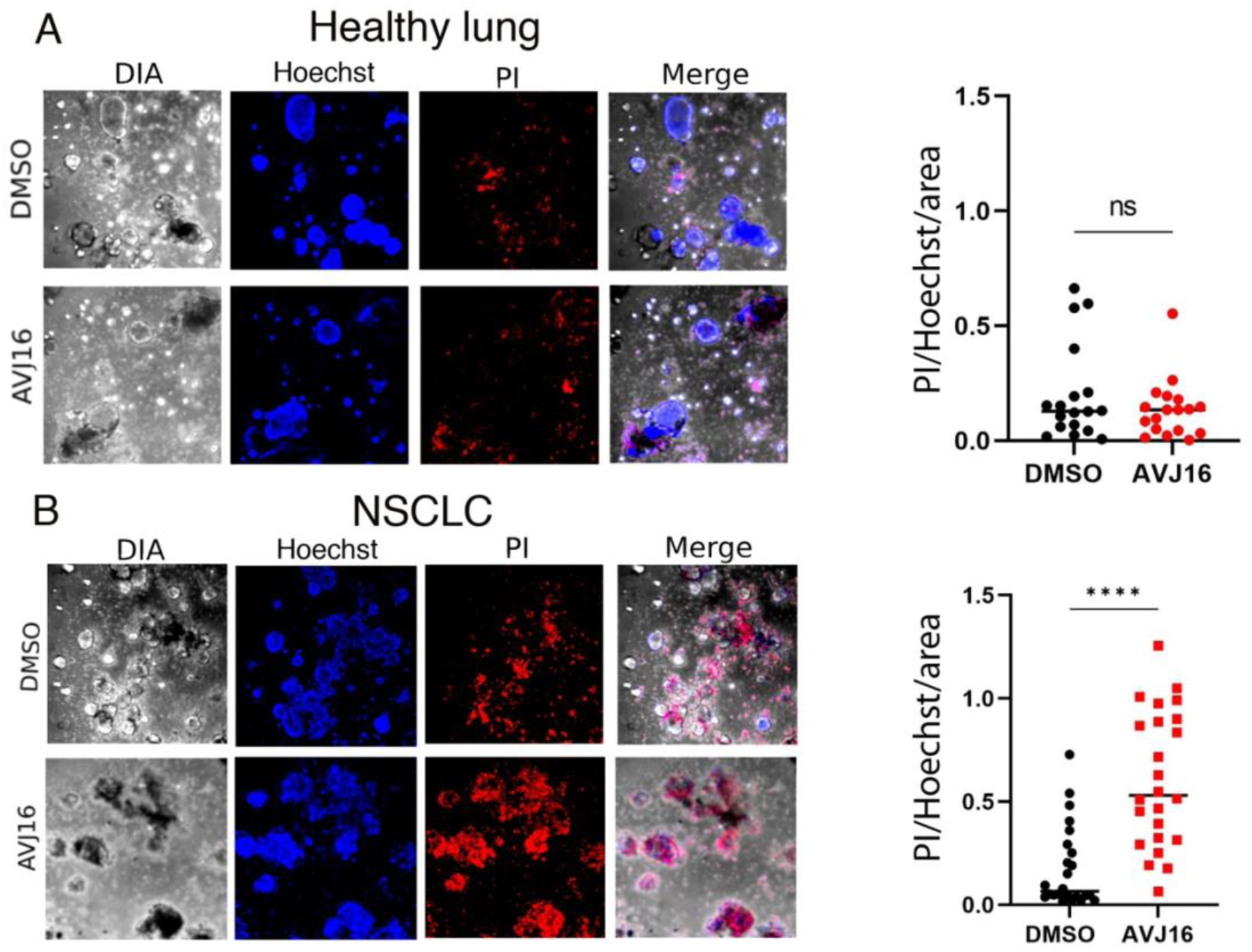
AVJ16 inhibits human NSCLC organoid growth. Organoids from normal human lungs (A) or from a human NSCLC lung cancer that expresses IGF2BP1 (B) were grown and treated with DMSO or AVJ16 for 72 hours, fixed, and then stained with Hoechst and propidium iodide (PI). The amount of PI staining per Hoechst staining per area was compared for each treatment and shown on the right. At least 18 organoids were analysed and measured per experiment. Biological duplicates were performed for each experiment.

## DISCUSSION

We have explored the mechanism of action of the IGF2BP1 inhibitor AVJ16 and examined its effects on lung carcinoma cells in 2D and 3D culture systems, and in mice. We have shown that AVJ16 engages IGF2BP1 in cells and induces genome-wide changes in the steady state levels of target RNAs and proteins. Many of these changes are associated with signaling pathways and cellular behaviors implicated in cancer phenotypes, and, accordingly, AVJ16 demonstrates anti-oncogenic effects in multiple assays. The specificity of AVJ16 is highlighted by our syngeneic mice model, where susceptibility to the inhibitor is dependent on the expression of the human IGF2BP1 gene. These results argue that AVJ16 could have potential as a targeted therapy for tumors expressing IGF2BP1.

Knowing that AVJ16 binds directly to IGF2BP1, both in vitro^14^ and in vivo (Fig. 1), we identified direct targets of IGF2BP1 in the presence or absence of the compound. eCLIP data show that almost ¾ of the over 22,000 peaks bound by IGF2BP1 are not affected by the compound. Mapping these peaks to genes reveals that a similar fraction of genes remains bound to IGF2BP1 even in the presence of the compound (4032/5181=78%). These results not only testify to the robust nature of the results but also emphasize the significance of those genes whose binding to IGF2BP1 is altered by AVJ16 treatment. Notably, most genes that contain a DBP contain only one DBP; and even in those that contain more than one, the regulation of the binding is highly coordinated (either all up or all down regulated). Thus, despite the fact that a gene may contain multiple binding sites, and even multiple DBPs, the coordinated regulation allows us to classify genes as upregulated or downregulated with respect to IGF2BP1 binding. The presence of DBPs and non-DBPs in the same gene further emphasizes the importance of site-specific regulation of IGF2BP1 binding sites and that their downregulation is not simply a result of downregulation of the gene itself.

Correlating the direct binding data with the RNAseq and proteomics data has provided valuable insights into the effects of AVJ16 treatment. Up or down regulation of a binding peak or peaks in a gene following AVJ16 treatment directly correlated with changes in the expression of the gene, as well as the protein encoded by that gene. This strongly argues that AVJ16 functions primarily by affecting RNA binding of IGF2BP1 to its target RNAs rather than through off-target RNAs or RBPs. Such off-target RNAs or RBPs would not be expected to correlate with the IGF2BP1 eCLIP data. These correlations also suggest that a major function of IGF2BP1 RNA binding in these cells is to stabilize the bound RNAs. Although this does not exclude additional roles for IGF2BP1, such as translational control or intracellular localization, these functions would be less likely to correlate with RNA steady state levels.

We have previously shown that AVJ16 appears to bind the hydrophobic groove between KH3 and KH4 in IGF2BP1, placing it in a position to disrupt RNA bound between the GXXR RNA binding loops in KH3 and KH4. Nevertheless, we show here that there are a significant number of RNAs whose binding is upregulated by AVJ16 treatment. A similar phenomenon was observed with the CuB inhibitor that binds KH12 in IGF2BP1, and it was suggested that this could be due to an allosteric change in the protein.^56^ Such an allosteric change in KH34 could also be induced by AVJ16. An additional, although not mutually exclusive, possibility is that disruption of an RNA’s binding at the KH34 site could allow for enhanced binding of that RNA at the KH12 site. Further investigation is necessary to determine how AVJ16 can affect both enhanced and reduced binding of IGF2BP1 to different RNAs.

Establishing a link between eCLIP-identified DBPs and changes in RNA/protein levels can potentially elucidate the effects of the inhibitor on cancer. For instance, the WNT signaling pathway was significantly impacted by AVJ16 treatment, with several WNT pathway genes identified by eCLIP as direct targets of IGF2BP1. AVJ16 treatment not only caused a reduction in the expression level of a number of WNT effector genes such as WNT5A and β-catenin but also an upregulation of a WNT inhibitor, DKK. Thus, the overall effect of AVJ16 treatment is a downregulation of WNT signaling. WNT signaling plays a critical role in regulating cell growth, differentiation, and survival during embryonic development and homeostasis. Aberrant activation of the WNT pathway is implicated in several types of cancer, including colorectal, breast, lung, and liver cancer. Upregulation of the WNT pathway can lead to the accumulation of β-catenin, which enters the nucleus and activates transcription of target genes. Moreover, the WNT pathway is also involved in the regulation of cancer stem cells, which are thought to contribute to tumor initiation, progression, and chemoresistance^57^. The importance of IGF2BP1 in regulating WNT target genes has recently been confirmed in CRC cells.^42^ The ability of AVJ16 to downregulate WNT activity can explain much of its success in inhibiting the pro-oncogenic behavior of LUAD cells in both 2D and 3D culture systems, as well as in mice xenografts.

Several studies have shown that upregulation of IGF2BP1 enhances chemoresistance, and downregulation of IGF2BP1 enhances chemosensitivity. shRNA-mediated knockdown of IGF2BP1 in melanoma, neuroblastoma, and CRC cell lines increases sensitivity to a variety of chemotherapeutic drugs, and IGF2BP1 expression is elevated in chemoresistant CRC tumors.^41,49,58,59^ Our RNAseq data indicate that several ABC transporters associated with multidrug resistance are downregulated by AVJ16 (including ABCG2^60^ and ABCC5^61^; Supplemental Data File 1). The fact that AVJ16 induces dye retention, often taken as a proxy for heightened chemosensitivity, suggests a potential use of AVJ16 as an adjuvant therapy to enhance effectiveness of other drugs. Indeed, genetic downregulation of IGF2BP1 sensitizes melanoma cells for both targeted therapy and immunotherapy^12,49^

AVJ16 is highly specific for the human IGF2BP1. Not only were para-tumoral injections effective at very low concentrations (0.05% of MTD) but also IP injections at 25% MTD were very effective at inhibiting distal xenografts that were already established. No detectable toxicity was observed at the concentrations used, as seen by the lack of inhibition of LKR-M explants not expressing human IGF2BP1.

The organ cultures showed a dramatic sensitivity to low doses of AVJ16, even stronger than the effects seen in the 3D culture systems described above. Notably, this particular tumor was taken from a patient that was LKB1 (STK11) negative. Although LKB1 is considered to be a tumor suppressor, NSCLCs with constitutively active Kras and inactivating mutations in LKB1 are highly sensitive to mTOR inhibitor.^62^ It will be of great interest to look for positive or negative factors that will enhance the effectiveness of AVJ16.

Taken together, the data presented here argue that AVJ16 can have a pronounced anti-cancer effect on lung carcinoma cells, both in vitro and in vivo, in a highly specific manner. These results suggest that AVJ16 may be useful as a targeted therapy for cancers expressing IGF2BP1.

## DATA AVAILABILITY

The eCLIP data have been deposited in NCBI’s Gene Expression Omnibus^63^ and are accessible through GEO Series accession number GSE274646 at https://www.ncbi.nlm.nih.gov/geo/query/acc.cgi?&acc=GSE274646. The RNAseq data have been deposited in NCBI’s Gene Expression Omnibus ^63^ and are accessible through GEO Series accession number GSE273982 at https://www.ncbi.nlm.nih.gov/geo/query/acc.cgi?acc=GSE273982. The mass spectrometry proteomics data have been deposited to the ProteomeXchange Consortium via the PRIDE^64^ partner repository with the dataset identifier PXD054641.

## ACKNOWLEDGEMENTS

We would like to express our appreciation to Professor Hannah Margalit for helpful discussions, Professor Zvi Fridlender and members of his lab for help with the xenograft experiments, the Interdepartmental Core Facilities of the Hebrew University Faculty of Medicine for help with the FACS analysis, fluorescence microscopy, RNAseq, and Incucyte analyses, Inbar Plaschkes and Yuval Nevo from the Bioinformatics unit at the Hebrew University Faculty of Medicine for their help with RNAseq analyses, Kylie Shen and the staff at Eclipse Bioinnovations for their help in analyzing the eCLIP data, and Hanan Schoffman at the Proteomic LC Mass Spectrometry Unit of the Hebrew University for help with the proteomics data.

## FUNDING

This work was supported by the Israel Science Foundation (JKY); the Israel Cancer Research Fund (JKY(; and Integra Holdings Ltd. (JKY).

## Notes

### Competing Interest Statement

The authors have declared no competing interest.

