## Supplemental Figures for "AVJ16 inhibits the RNA binding protein IGF2BP1 in lung adenocarcinomas and prevents tumor growth in mice"

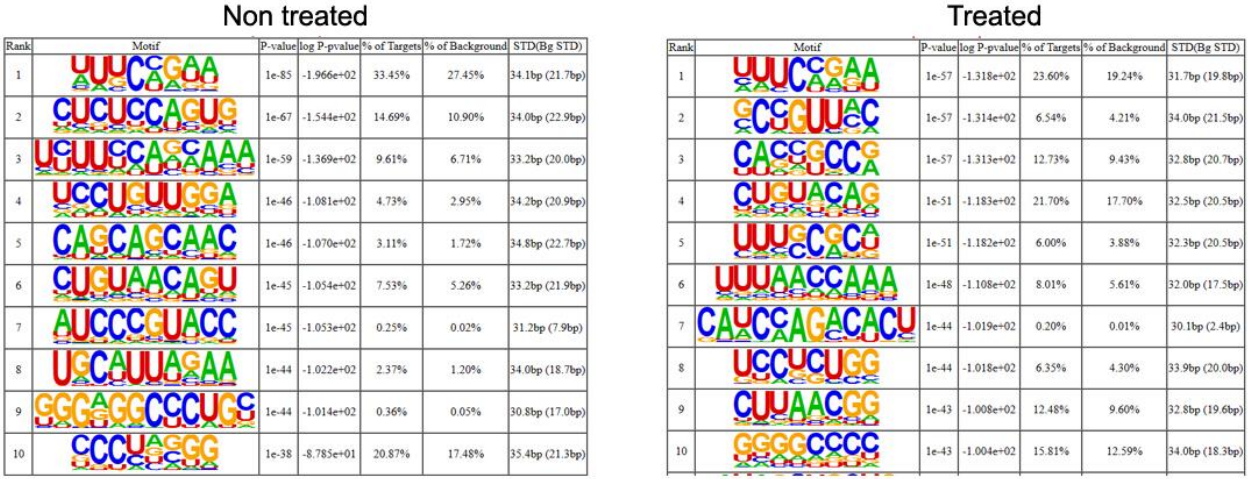

Non treated

| Rank | Motif | P-value | log P-pvalue | % of Targets | % of Background | STD(Bg STD) |
|---|---|---|---|---|---|---|
| 1 | UUUCCGAA | 1e-85 | -1.966e+02 | 33.45% | 27.45% | 34.1bp (21.7bp) |
| 2 | CUCUCCAGUG | 1e-67 | -1.544e+02 | 14.69% | 10.90% | 34.0bp (22.9bp) |
| 3 | UCUUCCAGCAAA | 1e-59 | -1.369e+02 | 9.61% | 6.71% | 33.2bp (20.0bp) |
| 4 | UCCUGUUGGA | 1e-46 | -1.081e+02 | 4.73% | 2.95% | 34.2bp (20.9bp) |
| 5 | CAGCAGCAAC | 1e-46 | -1.070e+02 | 3.11% | 1.72% | 34.8bp (22.7bp) |
| 6 | CUGUAACAGU | 1e-45 | -1.054e+02 | 7.53% | 5.26% | 33.2bp (21.9bp) |
| 7 | AUCCCGUACC | 1e-45 | -1.053e+02 | 0.25% | 0.02% | 31.2bp (7.9bp) |
| 8 | UGCAUUAGAA | 1e-44 | -1.022e+02 | 2.37% | 1.20% | 34.0bp (18.7bp) |
| 9 | GGGAGGCCCUGC | 1e-44 | -1.014e+02 | 0.36% | 0.05% | 30.8bp (17.0bp) |
| 10 | CCCUAGGG | 1e-38 | -8.785e+01 | 20.87% | 17.48% | 35.4bp (21.3bp) |

Treated

| Rank | Motif | P-value | log P-pvalue | % of Targets | % of Background | STD(Bg STD) |
|---|---|---|---|---|---|---|
| 1 | UUUCCGAA | 1e-57 | -1.318e+02 | 23.60% | 19.24% | 31.7bp (19.8bp) |
| 2 | GCCGUUAC | 1e-57 | -1.314e+02 | 6.54% | 4.21% | 34.0bp (21.5bp) |
| 3 | CACCGCCG | 1e-57 | -1.313e+02 | 12.73% | 9.43% | 32.8bp (20.7bp) |
| 4 | CUGUACAG | 1e-51 | -1.183e+02 | 21.70% | 17.70% | 32.5bp (20.5bp) |
| 5 | UUUGCGCA | 1e-51 | -1.182e+02 | 6.00% | 3.88% | 32.3bp (20.5bp) |
| 6 | UUUAACCAAA | 1e-48 | -1.108e+02 | 8.01% | 5.61% | 32.0bp (17.5bp) |
| 7 | CAUCCAGACACU | 1e-44 | -1.019e+02 | 0.20% | 0.01% | 30.1bp (2.4bp) |
| 8 | UCCUCUGG | 1e-44 | -1.018e+02 | 6.35% | 4.30% | 33.9bp (20.0bp) |
| 9 | CUUAACGG | 1e-43 | -1.008e+02 | 12.48% | 9.60% | 32.8bp (19.6bp) |
| 10 | GGGGCCCC | 1e-43 | -1.004e+02 | 15.81% | 12.59% | 34.0bp (18.3bp) |

**Supplemental Figure 1**. HOMER analysis of enriched motifs in eCLIP peaks from DMSO (Non-treated) and AVJ16-treated (Treated) H1299 cells.

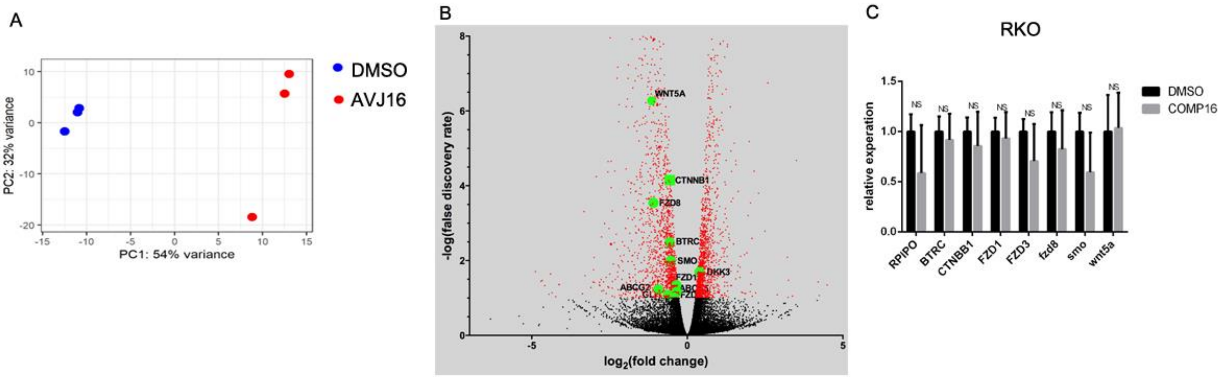


**Supplemental Figure 2.** <u>Impact of AVJ16 treatment on the steady state level of RNAs in H1299 cells</u>. RNA-seq was performed on RNA from cells treated with either 1.5µM of AVJ16 or DMSO for 48 hours. (A) Principal component analysis (PCA) plot showing segregation of AVJ16-treated and non-treated cells into two different groups (PC1 54%, PC2 32%). (B) Volcano plot representation of genes after AVJ16 treatment, highlighting some of the known targets of IGF2BP1, including βTrCP1, GLI1, and ABC transporters as well as the RNAs validated by qPCR in Figure 3E. (C) All seven WNT-related genes that were significantly downregulated in H1299 cells upon AVJ16 treatment were not significantly changed in abundance when assayed by qPCR in RKO cells, which do not express IGF2BP1.

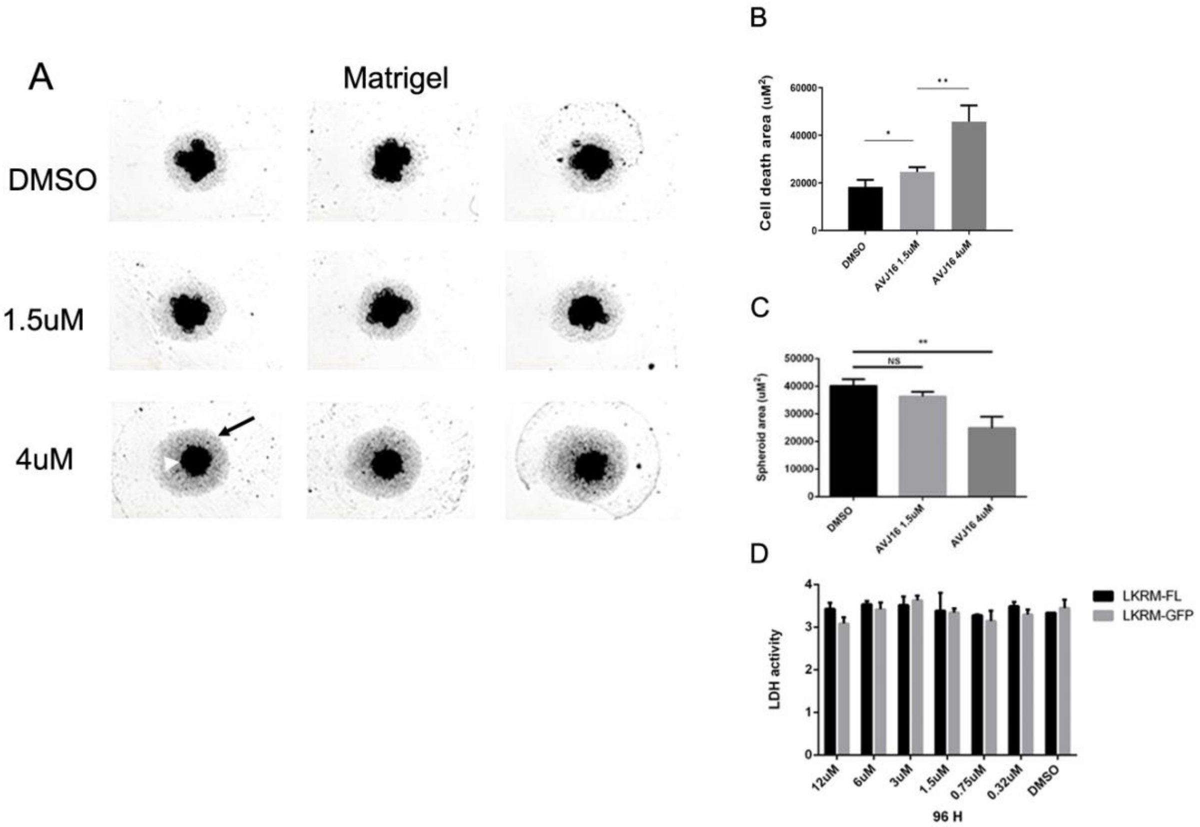


**Supplemental Figure 3**. <u>AVJ16 impeded growth and invasion capacity of H1299 cells grown as spheroids in the absence of Matrigel.</u> (A) Spheroids were grown in Matrigel in the presence of DMSO, 1.5μM AVJ16, or 4μM AVJ16. The white arrowhead shows an example of the live cells in a spheroid, and the black arrow points to the dead cells in the same sample. (B) Graph of the area of dead cells surrounding the spheroid as a function of treatment. (C) Graph of the viable cell area as a function of treatment. (D) Toxicity was measured by the presence of LDH in the medium after 96 hours of treatment at the indicated concentrations of AVJ16.

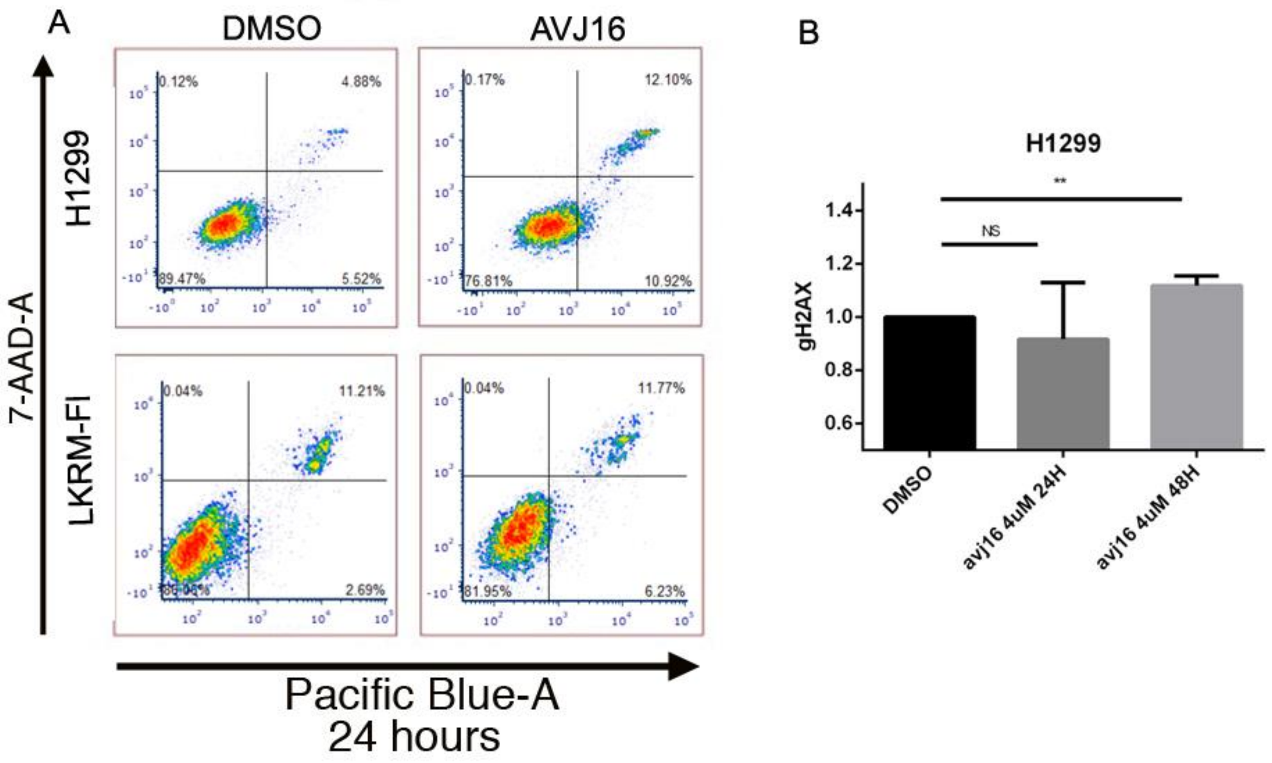


**Supplemental Figure 4.** <u>AVJ16 induces apoptosis</u> (A) FACS sorting was used to detect Annexin/7-AAD-A staining of H1299 and LKR-M-Fl cells treated with DMSO or 4μM AVJ16 for 24 hours. Late apoptotic cells appear in the top right quadrant. (B) H1299 cells treated with 4μM AVJ16 for 24 or 48 hours, and γ-H2AX levels, indicative of double strand DNA breaks, were measured by western blot analysis and normalized to a DMSO control.

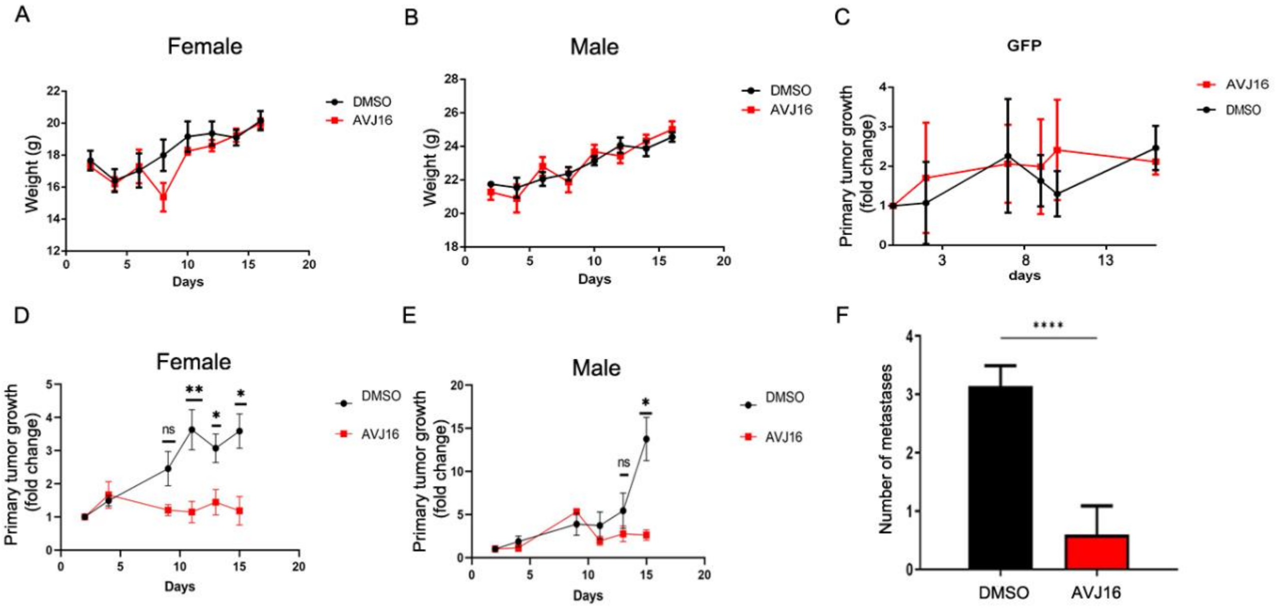


**Supplemental Figure 5.** AVJ16 effect on mice xenograft model. The female (A) and male (B) mice from the xenograft experiment described in Figure 8 were weighed over the course of the experiment from the point at which AVJ16 injections were begun (from day 12). (C) The effect of IP injections of AVJ16 or DMSO on the growth of LKR-M-GFP xenografts was monitored as in Figure 8. (D-F) Syngeneic mice were subcutaneously implanted with LKR-M-Fl cells and allowed to grow for 12 days, until the site of implantation was visible as a bulge. AVJ16 was injected peritumorally (PT) at the times indicated in Figure 8A. The primary tumor fold increase was measured every two days (time 0 - 12 days after subcutaneous injection of cells) in both females (D) and males (E). (F) The number of lung metastases was counted from lung histology specimens, of mice treated peritumorally (N=3).
